# A Kalirin Missense Mutation Enhances Dendritic RhoA Signaling and Leads to Regression of Cortical Dendritic Arbors Across Development

**DOI:** 10.1101/2021.03.22.436528

**Authors:** MJ Grubisha, T Sun, SL Erickson, L Eisenman, S Chou, CD Helmer, MT Trudgen, Y Ding, GE Homanics, P Penzes, ZP Wills, RA Sweet

## Abstract

Normally, dendritic size is established prior to adolescence then remains relatively constant into adulthood due to a homeostatic balance between growth and retraction pathways. However, schizophrenia is characterized by accelerated reductions of cerebral cortex gray matter volume and onset of clinical symptoms during adolescence, with reductions in layer 3 pyramidal neuron dendritic length, complexity, and spine density identified in multiple cortical regions postmortem. Nogo receptor 1 (NGR1) activation of the GTPase RhoA is a major pathway restricting dendritic growth in the cerebral cortex. We show that the NGR1 pathway is stimulated by OMGp and requires the Rho guanine nucleotide exchange factor, Kalirin-9 (KAL9). Using a genetically encoded RhoA sensor, we demonstrate that a naturally occurring missense mutation in *Kalrn*, KAL-PT, that was identified in a schizophrenia cohort, confers enhanced RhoA activitation in neuronal dendrites compared to wildtype KAL. In mice containing this missense mutation at the endogenous locus there is an adolescent-onset reduction in dendritic length and complexity of layer 3 pyramidal neurons in the primary auditory cortex. Tissue density of dendritic spines was also reduced. Early adult mice with these structural deficts exhibited impaired detection of short gap durations. These findings provide a neuropsychiatric model of disease capturing how a mild genetic vulnerability may interact with normal developmental processes such that pathology only emerges around adolescence. This interplay between genetic susceptibility and normal adolescent development, both of which possess inherent individual variability, may contribute to heterogeneity seen in phenotypes in human neuropsychiatric disease.

**SIGNIFICANCE STATEMENT:** Dendrites are long branching processes on neurons that contain small processes called spines that are the site of connections with other neurons, establishing cortical circuitry. Dendrites have long been considered stable structures, with rapid growth prior to adolescence followed by maintenance of size into adulthood. However, schizophrenia is characterized by accelerated reductions of cortical gray matter volume and onset of clinical symptoms during adolescence, with reductions in dendritic length present when examined after death. We show that dendrites retain the capacity for regression, and that a mild genetic vulnerability in a regression pathway leads to onset of structural impairments in previously formed dendrites across adolescence. This suggests that targeting specific regression pathways could potentially lead to new therapeutics for schizophrenia.

## INTRODUCTION

Schizophrenia (SZ) is a debilitating disease that affects approximately 1% of the population(1). Clinical symptoms, such as auditory hallucinations and delusions, emerge during the 2^nd^ or 3^rd^ decade of life. Recent studies, however, have emphasized that it is impairments in cognitive and sensory processes underlying the clinical symptoms that are the greatest contributors to functional impairment and long term disability(2-4). Specifically, auditory processing deficits have been consistently described at the neurophysiological level and include increased threshold for detecting differences between successive auditory stimuli and decreased amplitude of the mismatch negativity response to silent gaps, processes which require an intact of the auditory cortex(5, 6). Current therapeutics have limited efficacy for cognitive and sensory function impairments, and confer substantial morbidity owing to undesired side effects, underscoring the need for therapeutics targeting underlying molecular mechanisms.

Pyramidal cells (PCs) represent the most abundant neuronal type in the cerebral cortex(7), and their integrity is essential to the cognitive and sensory processes that are disrupted in SZ(5, 8). Among the most consistent and highly replicated findings from human postmortem studies of SZ are reductions in dendrite length, branching, and spine density in layer 3 PCs(9). Interestingly, these deficits appear to be more reliably defined in layer 3, as Layer 5 PCs have not been consistently shown to be impaired in postmortem studies of SZ(10). Because dendritic spines are the site of most excitatory synapses, much research to date has aimed to determine mechanisms of their reduction in SZ. However, total spine number is a function of both total dendrite length and spine density. Moreover, dendritic length and branching determine a PC’s receptive field(11, 12), help to segment computational compartments(13), and contribute substantially to how the received signals are integrated and transmitted to the cell body(14, 15). Thus, there is a compelling need to investigate dendritic length and branching alterations in SZ.

One of the driving physiological requirements for dendritic morphogenesis is the flexibility for adjustment in development and in response to experience(16). Initial rapid growth establishes a nearly full-sized dendritic arbor prior to adolescence(17). While dendrites retain the physiological flexibility to undergo modest changes in branch points or angles to refine circuitry, the overall net arbor size remains relatively constant across adolescence and into adulthood due to a homeostatic balance between growth and retraction pathways(17, 18).

Although rapid dendritic growth is complete prior to adolescence, this developmental epoch is a particularly active period of structural changes in the brain, leading to loss of cortical gray matter volume(19-22). In SZ, this reduction in gray matter volume is accelerated(23, 24), coincident with the onset of clinical symptoms(25, 26). The predominant component of cortical gray matter is neuropil, which comprises dendrites and axonal processes(27). It is possible that accelerated gray matter reductions during adolescence in SZ may be, in part, due to regression of dendritic architecture beginning during that time. It stands to reason, then, that a genetic susceptibility in a pathway involved in dendritic morphogenesis may be further exacerbated during the adolescent transition and lead to the onset of clinical symptoms of SZ.

Among the genes found to influence dendritic morphogenesis(16, 28) is *Kalrn*. Of the multiple isoforms generated from the *Kalrn* gene through alternative splicing, the longer isoforms (KAL9 and KAL12) possess 2 guanine nucleotide exchange factor (GEF) domains, the second of which activates the GTPase RhoA. A missense mutation (rs143835330) coding for a proline to threonine amino acid change in the *Kalrn* gene (*Kalrn-PT*) was first identified in a re-sequencing analysis in individuals with SZ(29). The P→ T change in *Kalrn-PT* is adjacent to the RhoA GEF domain in the KAL9 and KAL12 isoforms(29) and was shown to act as a modest gain-of-function for RhoA activity in a heterologous overexpression system(30). The activity of the first GEF domain, which activates Rac1, was shown to be unaltered by the *Kalrn*-PT mutation(30).

Importantly, RhoA regulation of dendritic morphogenesis requires molecular precision. Although constitutively active RhoA reduces dendritic morphogenesis in rodent PCs(31, 32), expression of dominant negative (DN) RhoA fails to affect dendritic outgrowth(33). Thus, targeting specific pathways upstream of RhoA activation is necessary to rescue structural impairments. For example, p75 is a neurotrophin receptor which directly binds to Nogo receptor (NGR) and, in response to ligand binding, subsequently activates RhoA(34, 35). In cerebellar granule neurons, the KAL9 isoform of the *Kalrn* gene has been shown to directly bind to p75 and provide the GEF domain required for RhoA activation(34). The NGR1/p75/KAL9 pathway is known to restrict neurite outgrowth(34). Specifically disrupting NGR1 mediated signaling via Nogo neutralizing antibodies promotes neurite outgrowth and extension *in vitro*(34, 36), and *in vivo* knockdown of neuronal-specific Nogo-A leads to increases in both branching and total length in L2/3 dendrites(37). Interestingly, increased levels of Nogo mRNA as well as elevated levels of KAL9 protein have been described in SZ(38, 39), suggesting enhanced activity of this pathway may contribute to the impairments in dendritic morphogenesis in disease. Although to date Nogo is the most well studied, numerous myelin-associated inhibitors (MAIs) have been identified as additional NGR1 ligands and similarly serve to limit neurite outgrowth(40). Interestingly, a highly potent MAI, oligodendrocyte-myelin glycoprotein (OMGp), increases in expression across adolescence(41).

Thus, we hypothesized that *Kalrn*-PT would act as a RhoA gain-of-function in neurons and lead to adolescent-onset reductions in dendritic length and complexity.

## RESULTS

### *Kalrn-PT* acts as a gain of function for RhoA signaling

Previous work from our lab demonstrated that expression of *Kalrn-PT* on the KAL9 background (KAL9-PT) increased RhoA activity compared to KAL9-WT in an *in vitro* heterologous overexpression system(42). However, primary neuronal culture is superior in that it captures features of normal neuronal development and endogenous kalirin expression, features lacking in heterologous systems. Thus, we derived cortical neuronal cultures from E18 rats and overexpressed either KAL9-WT or KAL9-PT at equivalent levels (p=0.26 between genotypes, SI appendix, Figure S1). At the time of transfection with the Kalirin-containing plasmids, neurons were also transfected with a FRET-based genetically encoded RhoA sensor (RhoA2G)(43). As previously described(44), regions of interest (ROIs; 5–10 μm boxes) were defined along the dendritic shaft and soma, affording the ability to determine RhoA activation with subcellular localization.

Representative neurons with the defined ROIs are shown in Figure 1 panels A-C, with activity represented as a heat map. While overexpression of KAL9-WT showed no increase baseline RhoA activity in dendrites compared to control, KAL9-PT increased RhoA activity relative to KAL9-WT (Fig 1D). This Kal9-PT increase in RhoA was more pronounced in mid-proximal dendritic shaft regions, corresponding to 30-50 µm from the soma (Fig 1E).

**Figure 1.**
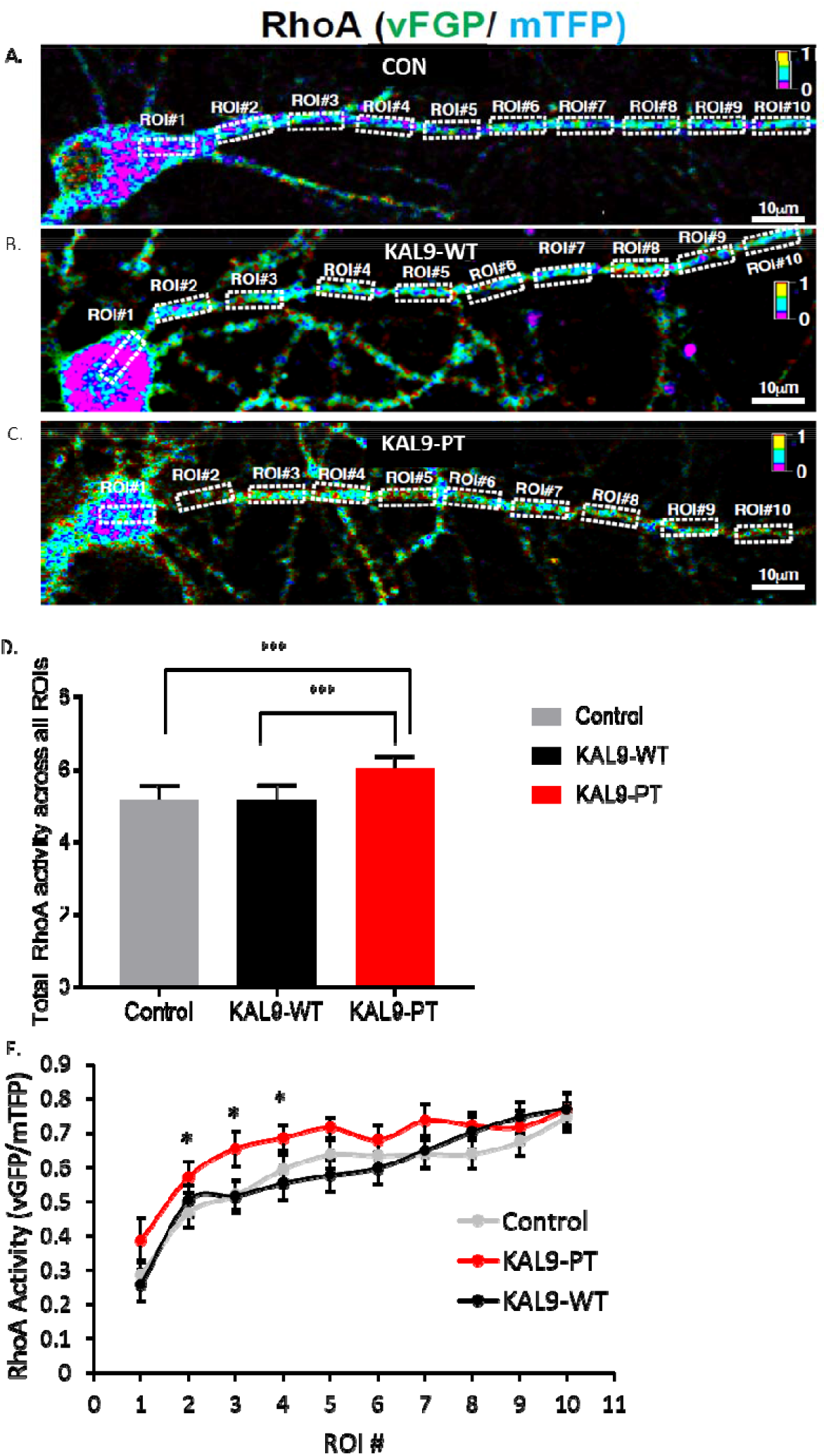
KAL-PT confers gain-of-function for RhoA activity in cortical neurons. (A) Representative image of RhoA sensor activity as measured by FRET imaging in control (A), KAL9-WT (B), and KAL9-PT (C) transfected neurons (transfected at DIV8, imaged at DIV14-18). ROIs are indicated by dashed yellow boxes, with 1 ROI within the soma and 2-10 defined along a dendritic shaft. Activity is represented as a heat map, with warmer colors indicating increased activity. (D) Sum all of RhoA activity across all ROIs demonstrates a significant increase in RhoA activity in KAL-PT compared to control and KAL-WT. (E) KAL9-PT shows increased RhoA sensor activity within proximal dendritic shaft ROIs (*p<.05 between KAL-WT and KAL-PT). N= 10 neurons per condition. Data shown are ± SEM. ***p<.0001

### OMGp signaling in pyramidal neurons limits dendritic arborization via an NGR1 and KAL9 dependent pathway

In cerebellar granule neurons, KAL9 mediates GEF activation of RhoA downstream of NGR/p75 signaling(34). This serves to limit early neurite outgrowth. We hypothesized that in excitatory pyramidal neurons, KAL9 might also regulate RhoA signaling downstream of NGR1 signaling via this same pathway. We further hypothesized that in pyramidal neurons this pathway may act on dendritic arbors. To test this, we first utilized hippocampal slice culture, as the role of NGR1 in reducing dendritic arbor size has been well characterized in this system(45). We overexpressed NGR1 in CA1 pyramidal cells and observed decreased dendritic length and complexity compared to GFP-only control, as previously described. Concomitant transfection with a previously validated shRNA directed at KAL9(34) rescued this phenotype, suggesting that KAL9 is required for NGR1 signaling to restrict dendritic growth (Fig 2).

**Figure 2.**
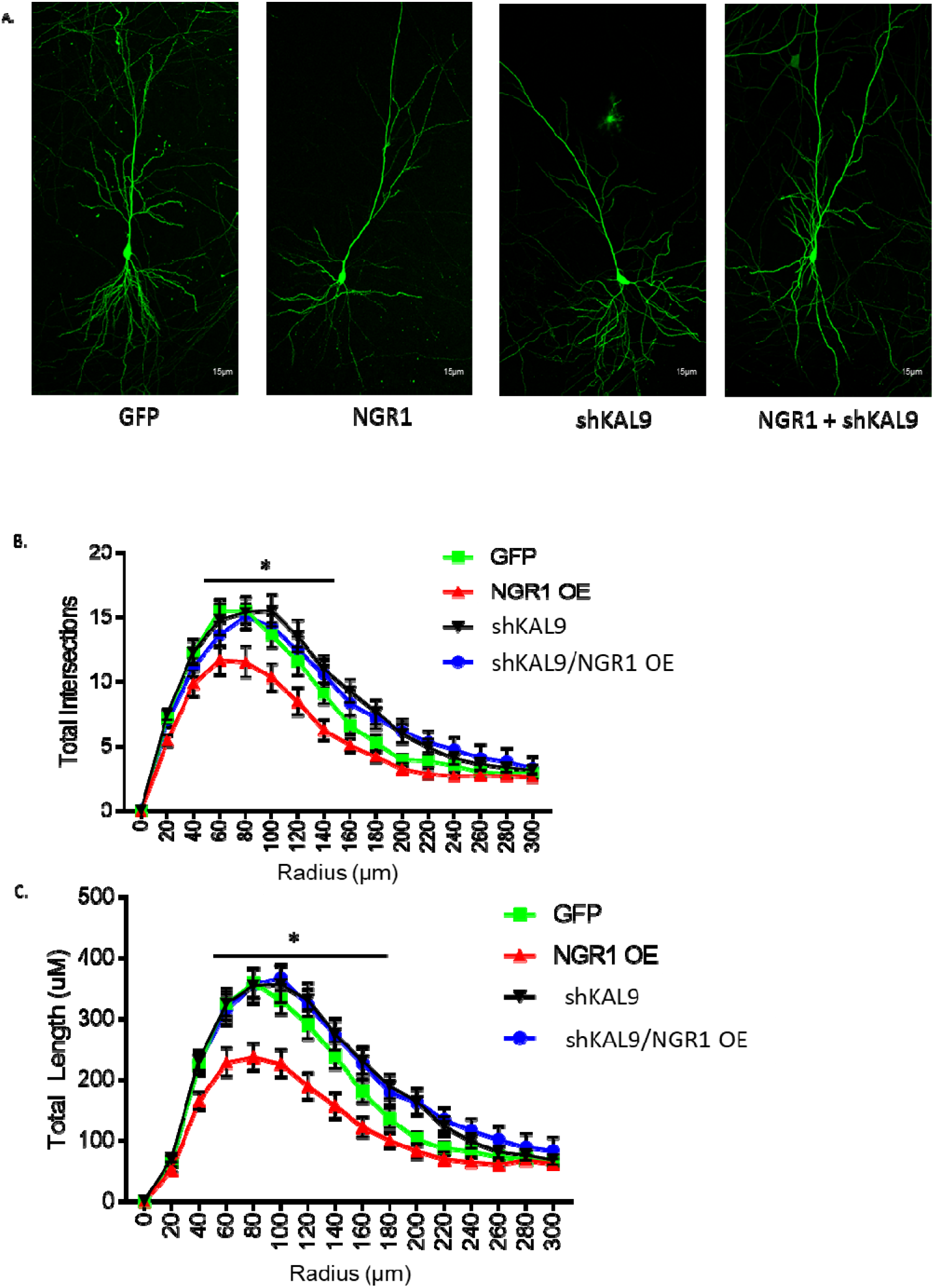
KAL9 is necessary for NGR1-mediated dendritic growth restriction. (A) Representative images of CA1 pyramidal neurons expressing GFP +/-NGR1 or shKAL9. (B) Sholl analysis of total intersections shows a decrease in dendritic complexity with NGR1 overexpression (*p=.03) that is rescued by concomitant transfection with shKAL9. (C) Sholl analysis of total length shows a decrease in dendritic length with NGR1 overexpression (*p=.01) that is rescued by concomitant transfection with shKAL9. N= 20-24 neurons per condition from 3 independent experiments. Data shown are ± SEM.

NGR1 is known to signal through multiple pathways, thus we conducted a second series of experiments to further define the role of KAL9 in mediating NGR1-dependent effects on dendritic architecture. To do this, we used the MAI, OMGp, which has previously been shown to act through NGR1 to inhibit early neurite outgrowth(46). However, OMGp has been observed to be abundantly expressed by adult CNS neurons where its role remains unknown(47). Thus, we hypothesized that OMGp would signal through the NGR1 and require KAL9 to restrict dendritic arborization in pyramidal neurons (Figure 3A). Treatment of dissociated cortical neurons (48h treatment from DIV12-14) with OMGp (200ng) leads to reductions in both length (p<0.0001) and complexity (p<0.0001) of dendritic arbors (Figure 3B,C), suggesting a potential role for this protein in regulating dendritic architecture. We next genetically ablated NGR1 using a previously validated shRNA construct(45) and found that knockdown of NGR1 expression mitigates the effect of OMGp on dendritic architecture (Fig 3D,E). This suggests OMGp functions via the NGR1 signaling pathway to inhibit dendrite growth. Finally, we genetically ablated KAL9 expression through use of a validated shKAL9 construct(34) and also observed OMGp failed to produce a significant effect on dendritic architecture in the absence of KAL9 (Fig 3F,G). Taken together, these findings demonstrate OMGp can restrict dendritic length and complexity in cortical neurons, and this effect requires both NGR1 and KAL9.

**Figure 3.**
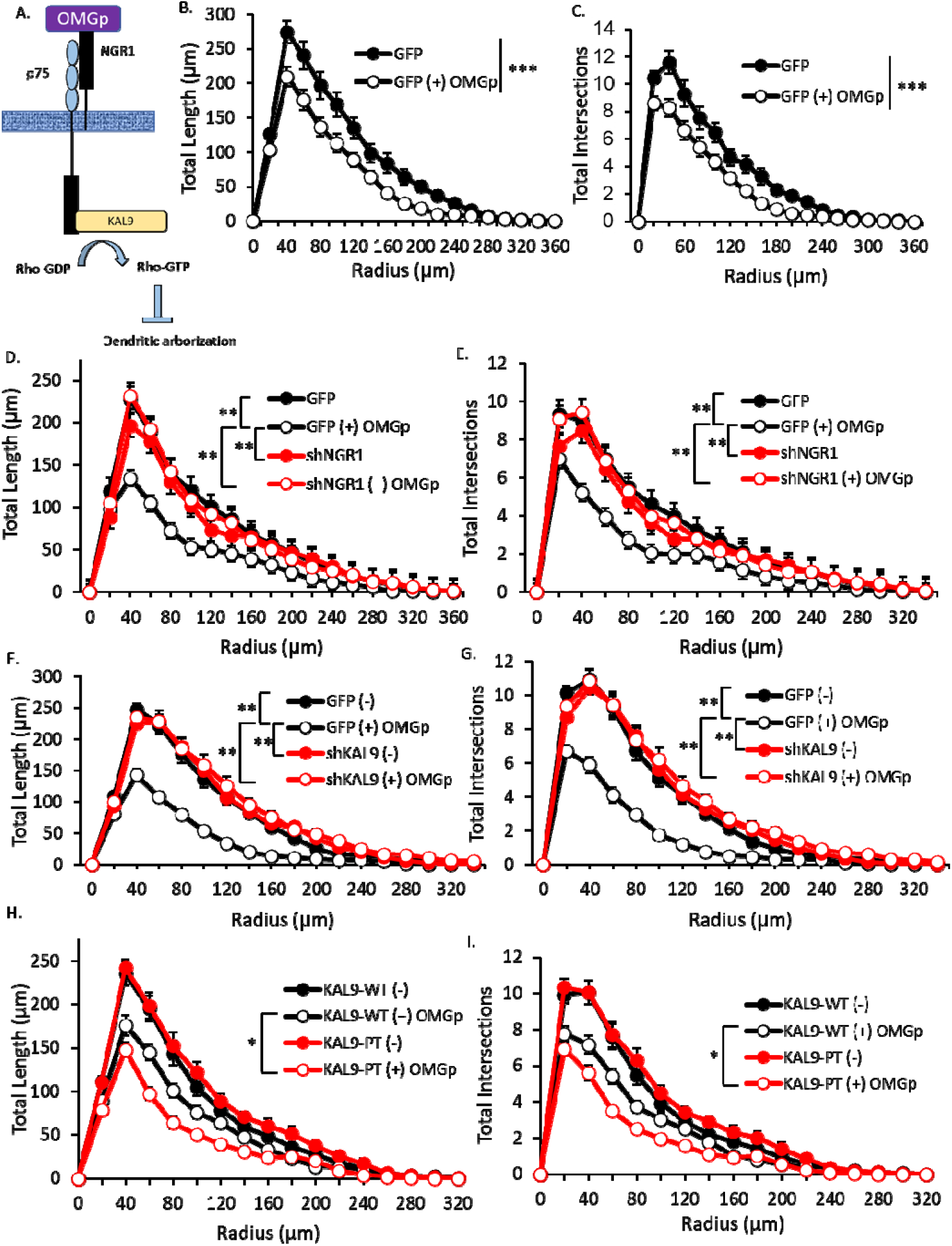
Dendritic growth restriction by OMGp requires both NGR1 and KAL9. (A) Schematic of the OMGp-NGR1/p75/KAL9 pathway that impedes dendritic arborization. (B) 48h OMGp (200ng) treatment of DIV12-14 cortical neurons reduces total dendritic length (C) and complexity. (D) Genetic ablation of NGR1 expression mitigates the effect of OMGp treatment on dendritic length (E) and complexity in cortical neurons. (F) Genetic ablation of KAL9 expression mitigates the effect of OMGp treatment on dendritic length (G) and complexity in cortical neurons. Overexpression of KAL9-PT shows further reductions in dendritic (H) length and (I) complexity in response to OMGp treatment compared to overexpression of KAL9-WT. N=26-30 neurons per condition from 3 independent experiments. Data shown are ± SEM. ***p<0.0001, **p<0.01, *p<0.05

Given the increased RhoA activity downstream of KAL9-PT, we next sought to determine if the effect of OMGp signaling on dendrites via NGR1/p75/KAL9 was enhanced by KAL9-PT compared to wild type. To test this, we transfected dissociated cortical cultures with either KAL9-WT or KAL9-PT on DIV8, then stimulated with either PBS alone or 200ng OMGp for 48h on DIV12-14. Neurons were fixed and dendritic arbors were reconstructed. As Figure 3H,I shows, OMGp significantly decreased both length and complexity of dendritic arbors in both KAL9 genotypes. This effect was more pronounced in the KAL9-PT neurons versus KAL9-WT (p=0.012 between KAL9-WT (+) OMGp and KAL9-PT (+) OMGp for dendritic length; p=0.01 between KAL9-WT (+) OMGp and KAL9-PT (+) OMGp for dendritic intersections), suggesting KAL9-PT enhances NGR1-RhoA signaling resulting in more pronounced dendritic growth deficits.

### *Kalrn-PT* mice as a model of the P→T missense mutation

To determine the impact of the *Kalrn*-PT mutation *in vivo* and preserve the physiologic stoichiometry of kalirin, we introduced the missense mutation at the endogenous locus of the C57BL/6J mouse strain using CRISPR/Cas9 gene editing technology. We designed a custom targeted single guide RNA (sgRNA94) and a repair oligonucleotide containing the C→A missense mutation constituting *Kalrn-PT* (see SI appendix, Table S1, Methods). In addition to the missense mutation, a silent mutation located in the 3^rd^ base pair of the codon harboring the missense mutation was introduced (C→T). This silent mutation produced a unique restriction site for Hinf1 (Fig 4A). Thus, restriction digest could be used for rapid and reliable genotyping assays once the line was established.

**Figure 4.**
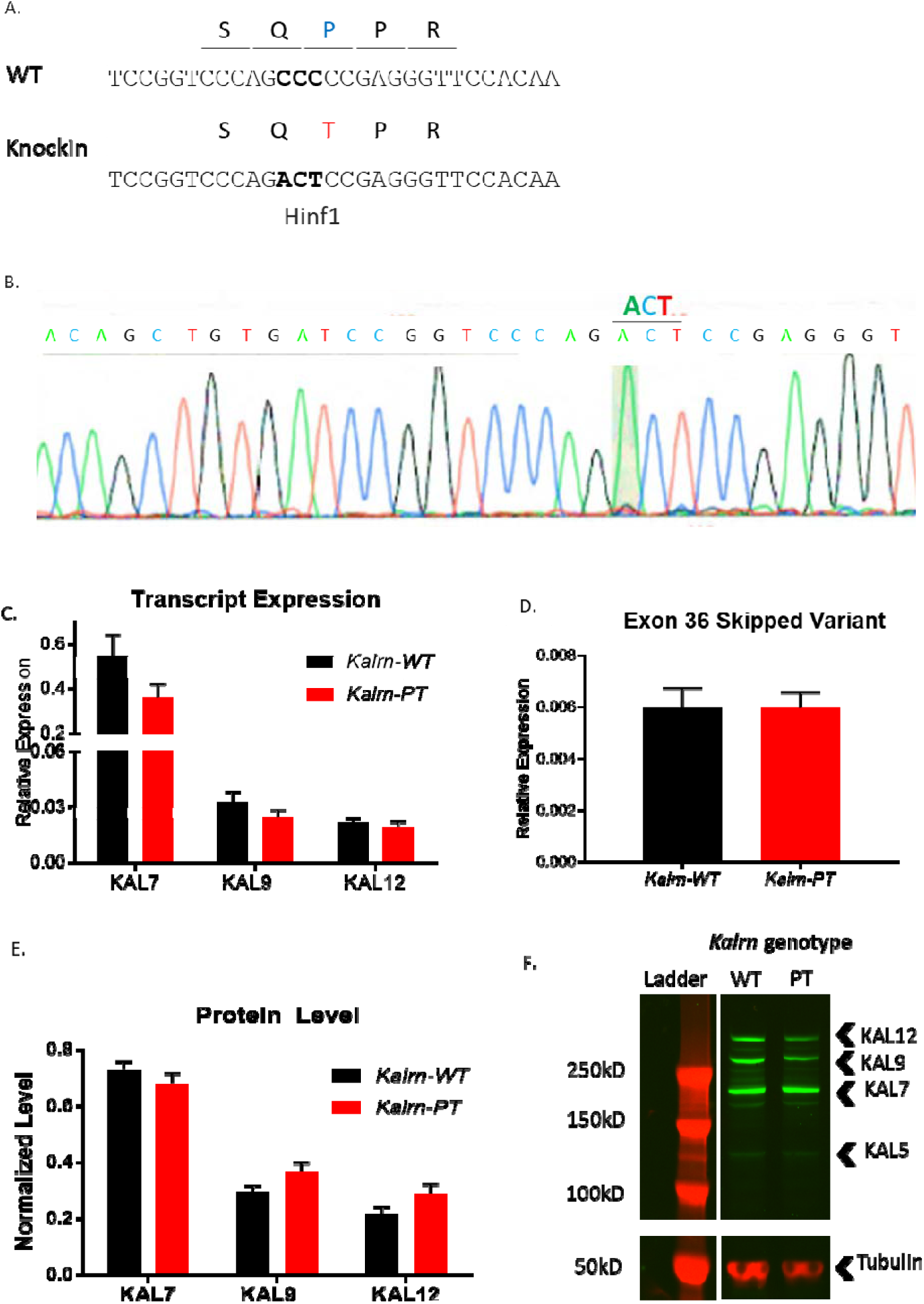
A genetic mouse model of *Kalrn*-PT was generated using CRISPR/Cas9. (A) Partial cDNA and protein sequences in wildtype and knockin mice. A silent mutation in the targeted codon introduces a unique Hinf1 restriction site. Targeted codon is in bold. (B) Confirmatory sequencing was performed on the founder and F1 generation, demonstrating homozygous knockin of the mutation. The site of the mutation is designated above the sequence. Gray matter homogenate from WT or *Kalrn-PT* homozygous mice (P28-P34, N=7 per group) was subjected to (C,D) RT-qPCR or (E,F) Western blotting to confirm transcript and protein expression levels of all KAL isoforms. No change in expression was observed as a result of genotype. There was no difference in exon skipping (exon 36) between genotypes. F is a representative blot of a pair of animals, *Kalrn*-WT on the left and *Kalrn*-PT on the right. All major isoforms are present in expected ratios at the given age, KAL5 is present in minor amounts and was not quantified. Data shown in panels C-F are from N=7 animals/genotype and are displayed ± SEM.

A founder male mouse who was homozygous for the *Kalrn-PT* mutation was chosen for line establishment. Confirmatory Sanger sequencing was performed to verify presence of both the missense and silent mutations in the correct region (Fig 4B). To confirm absence of any unintended off-target effects, we used prediction software to generate 18 sites with high homology to our guide sequence. Our founder mouse was confirmed to be free of off target effects at all sites tested by both PCR and Sanger sequencing (see SI appendix, Table S2, Methods). To establish the *Kalrn-PT* line, we bred our founder mouse with a purchased wildtype female of the C57BL/6J strain. Although the founder mouse was confirmed to be homozygous for the *Kalrn-PT* mutation, this was assessed only in peripheral cells derived from a tail snip sample and did not account for any mosaicism that exists in the gametes. Knockin mice are known to possess mosaicism (48), thus we monitored the rate of mutation transmission in the F1 generation as well as performed Sanger sequencing to confirm genotypes in all F1 generation derived from the founder mouse. It was determined that our founder mouse was transmitting the *Kalrn-PT* mutation with approximately 50% efficiency, and all F1 offspring retained for the line were confirmed to have only the missense and silent mutations at the desired locus. Subsequent generations were determined to be transmitting the mutation at expected Mendelian ratios.

We next evaluated whether the *Kalrn-PT* mutation was associated with altered levels of *Kalrn* mRNA and protein. Kalirin isoform expression is known to be developmentally dependent, with isoforms 9 and 12 being the predominate isoforms for the first 2 weeks of age before kalirin-7 expression increases and becomes the dominant isoform (49, 50). Our assays were performed on P28-35 animals, an age at which Kalirin-7 would be expected to be the most abundant isoform. We performed quantitative real-time PCR using primers specific for the 3 most abundant kalirin isoforms, specifically Kalirin-7,-9, and-12. There was no influence of genotype on the expression levels of any of the major Kalirin isoforms. Given recent data that demonstrate an increased frequency of exon skipping in Kalirin (exon 36) associated with schizophrenia (51), we also examined if there was any impact of the *Kalrn-PT* mutation on increased frequency of this naturally-occurring splice variant. As Figure 4D shows, there was no change in the relative expression of this variant as a result of genotype. Importantly, there was no change in transcript expression levels of the affected isoforms (i.e. Kalirin-9 and −12) and there was no compensatory change in expression levels of Kalirin-7 (Fig 4C). We further confirmed endogenous levels by evaluating the protein expression by western blot. Again, there was no change in levels of any of the major Kalirin isoforms as a result of genotype (Fig 4E,F).

### Dendritic morphogenesis is impaired in *Kalrn-PT* mice *in vivo*

Abnormalities in sensory processing (5) as well as reductions in PC somal size (52), spine density (53), and neuropil (54) have all been described in schizophrenia. These functional and structural deficits implicate impairments in Layer 3 of primary auditory cortex (A1) (8). Thus, we initially chose to focus on Layer 3 PCs in A1 to test the effects of *Kalrn-PT* on dendritic morphology. We generated a cohort of 12-week ± 3 days mice and used Golgi-Cox staining to detail neuronal structure. A total of 5 animals/genotype/sex were included and vaginal cytology on female animals was collected at time of sacrifice. Based on pilot studies, the variabilities across neurons from the same layer and region within animal were found to be quite small. We thus chose six neurons per animal for imaging and reconstruction. While our sample size was underpowered to detect primary effects of sex, our groups were balanced for sex across genotypes, ensuring that we did not introduce systematic bias. We did nonetheless test the effect of sex in our small sample size to ensure that there was no significant effect of sex on any of the dendritic measures in our cohort (SI appendix, Table S3), and thus the groups were combined to provide a total of 10 wildtype and 10 *Kalrn-PT* homozygous animals. Using Sholl analysis, we found significant reductions in dendritic length and complexity across both the apical (Fig 5C, D) and basilar (Fig 5E, F) arbors (p=0.0002 and p<0.0001, respectively). Representative reconstructions are shown in Fig 5A, B. In contrast to layer 3 PCs, those found in layer 5 have not been consistently shown to be impaired in postmortem studies of SZ(10). Using the same approach as described above, we found that there was no significant difference in dendritic length or complexity in layer 5 PCs between genotypes (SI appendix, Fig S3).

**Figure 5.**
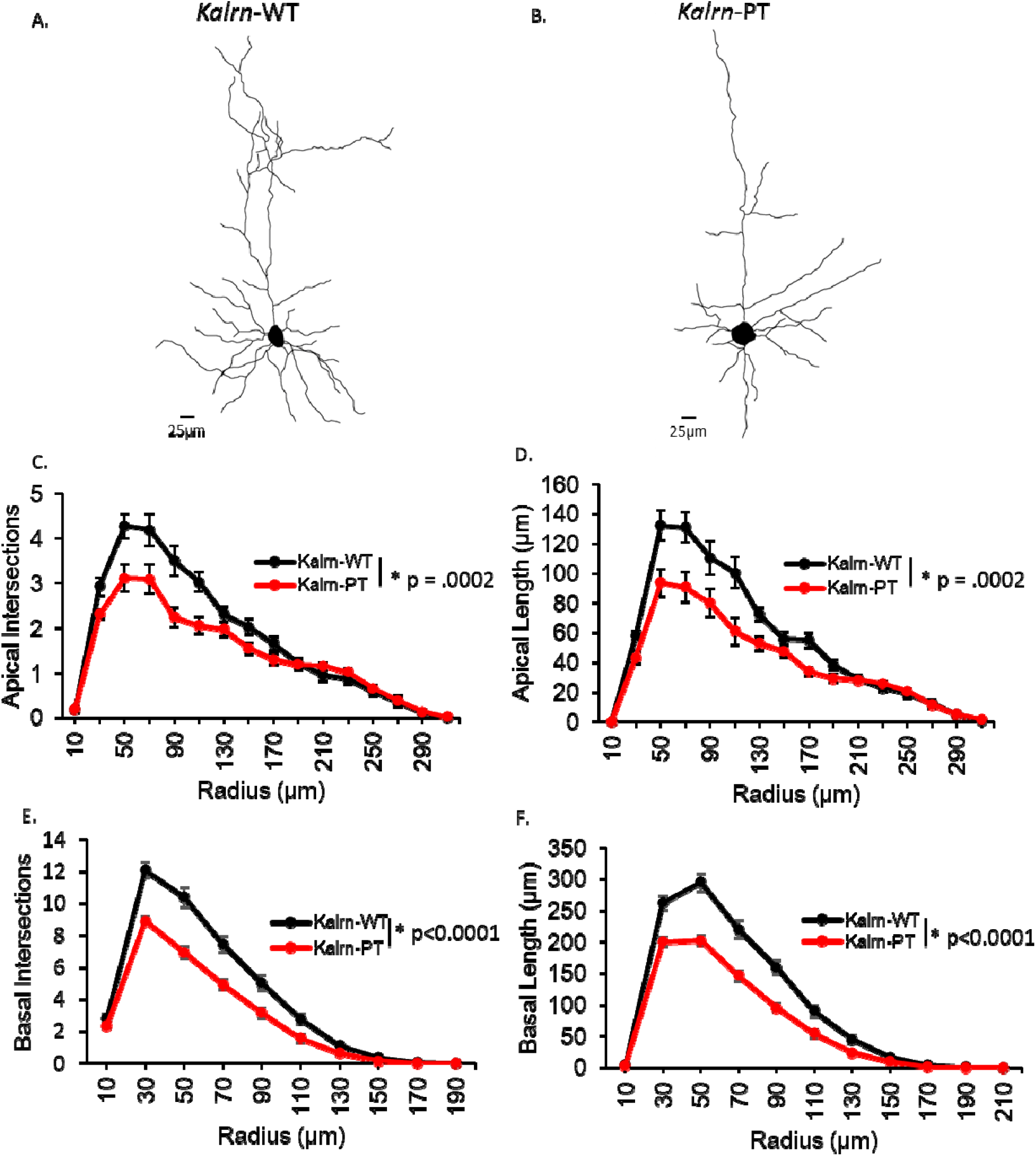
*Kalrn*-PT leads to reductions in dendritic architecture at 12-weeks. Dendritic reconstructions from Golgi stained PCs in A1 of 12-week old (A) *Kalrn*-WT and (B) *Kalrn-PT* mice demonstrate reduced dendritic complexity and length in (C, D) apical arbors as well as (E, F) basilar arbors. (N=10 mice per genotype, 6 neurons per animal) Results shown are from repeated measures ANOVA with main effect of genotype. p-values for each analysis are indicated within the corresponding graph. Data shown are average values across animals ± SEM.

In addition to reductions in dendritic length and complexity in L3 PCs in schizophrenia, human postmortem studies consistently demonstrate reduced L3 dendritic spine number and spine density per unit of tissue in A1 in schizophrenia (9, 55). To determine if dendritic spine density was also altered in *Kalrn-PT* mice, we used the Golgi material generated from our 12-week cohort described above. We added an additional animal per genotype in order to balance estrus phase across female animals (2 animals/phase/genotype from 3 phases: estrus, metestrus, proestrus; see SI appendix, Table S3a). Balancing estrus phase across groups ensured we did not introduce any systematic bias, and previous work in rodent cortex has shown that steady state cortical spine density does not differ across estrus phase(56). Nonetheless, we did evaluate estrus phase as a potential covariate despite our small sample size, and found no significant effect (Si appendix, Table S3b), consistent with the literature. Figure 6A,B shows a representative image of a PC containing a secondary apical dendrite along which spines were manually counted, along with representative images from *Kalrn-WT* (top panel) and *Kalrn-PT* (bottom panel) mice. Quantitative data are shown in Panel C, demonstrating no mean difference in spine density per unit dendrite length across genotypes (p=0.73). However, spine number and spine density per unit of tissue are a function of both total dendritic length and spine density per unit length. To estimate spine tissue density, we used spinophilin/phalloidin colocalization(9). Both spinophilin and phalloidin localize to dendritic spines(57-59), and previous work has validated the co-localization approach as a method for identifying presumptive spine(9). As shown in Fig 6D,E, we found a reduction in spine tissue density in *Kalrn*-PT mice compared to *Kalrn*-WT (p=.05). Furthermore, given the reductions in dendritic length and complexity, we also tested if total gray matter volume reductions were present. Volume measurements performed on the neocortex of a small cohort of 12-week old mice showed a 4% reduction in gray matter volume, though this did not achieve statistical significance (p=0.24, SI appendix, Fig S3).

**Figure 6.**
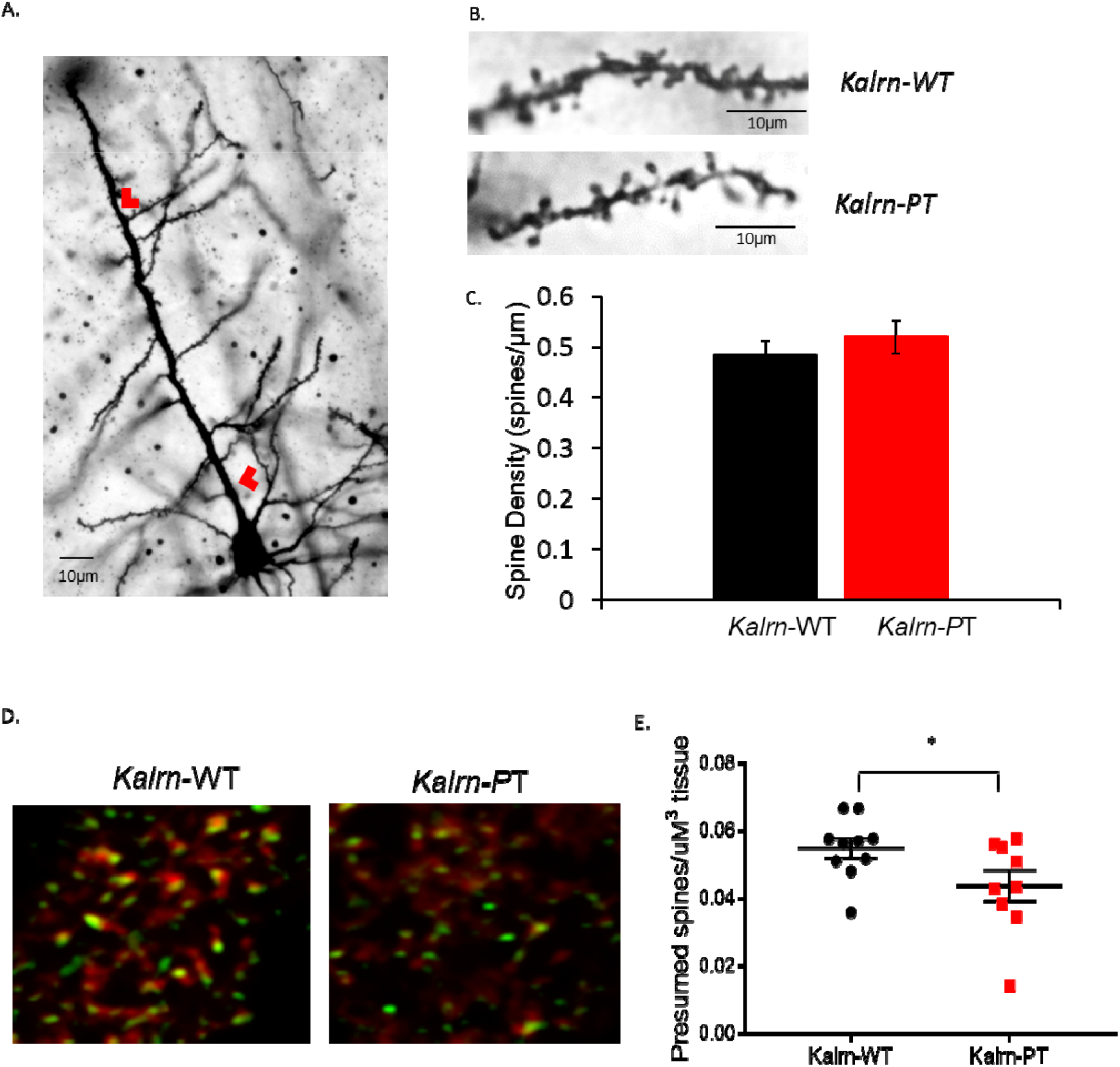
*Kalrn*-PT mice have reduced spine number compared to *Kalrn*-WT. (A) Representative Golgi image of a PC imaged for spine density analysis. Red arrowheads indicate designated length long primary apical dendrite wherein any secondary arising within that area was included for density analysis. (B) Representative images of dendritic segments on which spines were counted *(Kalrn-WT* image shows 0.49 spines/µm, *Kalrn-PT* image shows 0.51 spines/µm) (C) No change in spine density was observed as a result of genotype (*Kalrn-WT* average 0.49 spines/µm, *Kalrn-PT* average 0.52 spines/µm; p=0.73; N=10 animals/genotype). (D) Co-localization of phalloidin-labeled (red) and spinophilin-immunoreactive (green) puncta in Layer 3 of A1 from *Kalrn-WT* and *Kalrn-PT* mice (N=10 animals per genotype) were used to indicate presumptive spines. Ten sites per animal were imaged. Representative images for each genotype are shown. (E) Data shown are number of spine objects/µm^3^ of tissue for each animal. There is a decrease in presumptive spine objects in the *Kalrn-PT* mice compared to WT (*p=.05) All data shown are ± SEM.

### *Kalrn*-PT mice exhibit increased gap duration threshold

To test if the structural deficits observed in auditory cortex L3 PCs were associated with functional impairments, we sought to test cortical auditory function in a 12-week cohort of wildtype and *Kalrn*-PT mice. We used the acoustic startle response paradigm in which the extent of response inhibition by a prepulse (prepulse inhibition, PPI) is quantified. When the prepulse is a noise at a lower sound level pressure than the startle-eliciting noise (noise-PPI), PPI is dependent upon subcortical auditory processing(60). Conversely, detection of silent gaps embedded in noise has been shown to require the primary auditory cortex(61, 62). If the animal detects the gap, the startle response is inhibited. Thus, use of a modified PPI paradigm in which the prepulse is a silent gap (Gap-PPI) can be used to assess the integrity of the auditory cortex.

We generated a cohort of male and female mice, aged 12 weeks (N=11 *Kalrn*-WT, N=12 *Kalrn*-PT) and tested them for baseline acoustic startle response, noise-PPI, and Gap-PPI. We established that there is no significant difference in baseline acoustic startle response between genotypes (Fig 7A; *Kalrn*-WT = 0.435 ± 0.049N, *Kalrn*-PT = 0.339 ± 0.041N, p=0.15). Noise-PPI is similarly equivalent between genotypes, suggesting that subcortical auditory processing is unaffected by the *Kalrn*-PT mutation (Fig 7B). To assess cortical auditory processing, we examined thresholds for Gap-PPI, defined as the shortest gap durations to elicit significant PPI. The threshold for Gap-PPI increased from 4ms in *Kalrn*-WT mice to 20ms in the *Kalrn*-PT mice (Fig 7C,D; thresholds indicated by arrows). Taken together, these results suggest that, compared with *Kalrn*-WT mice, the *Kalrn*-PT mice exhibit decreased detection of gap-in-noise at shorter gap durations.

**Figure 7.**
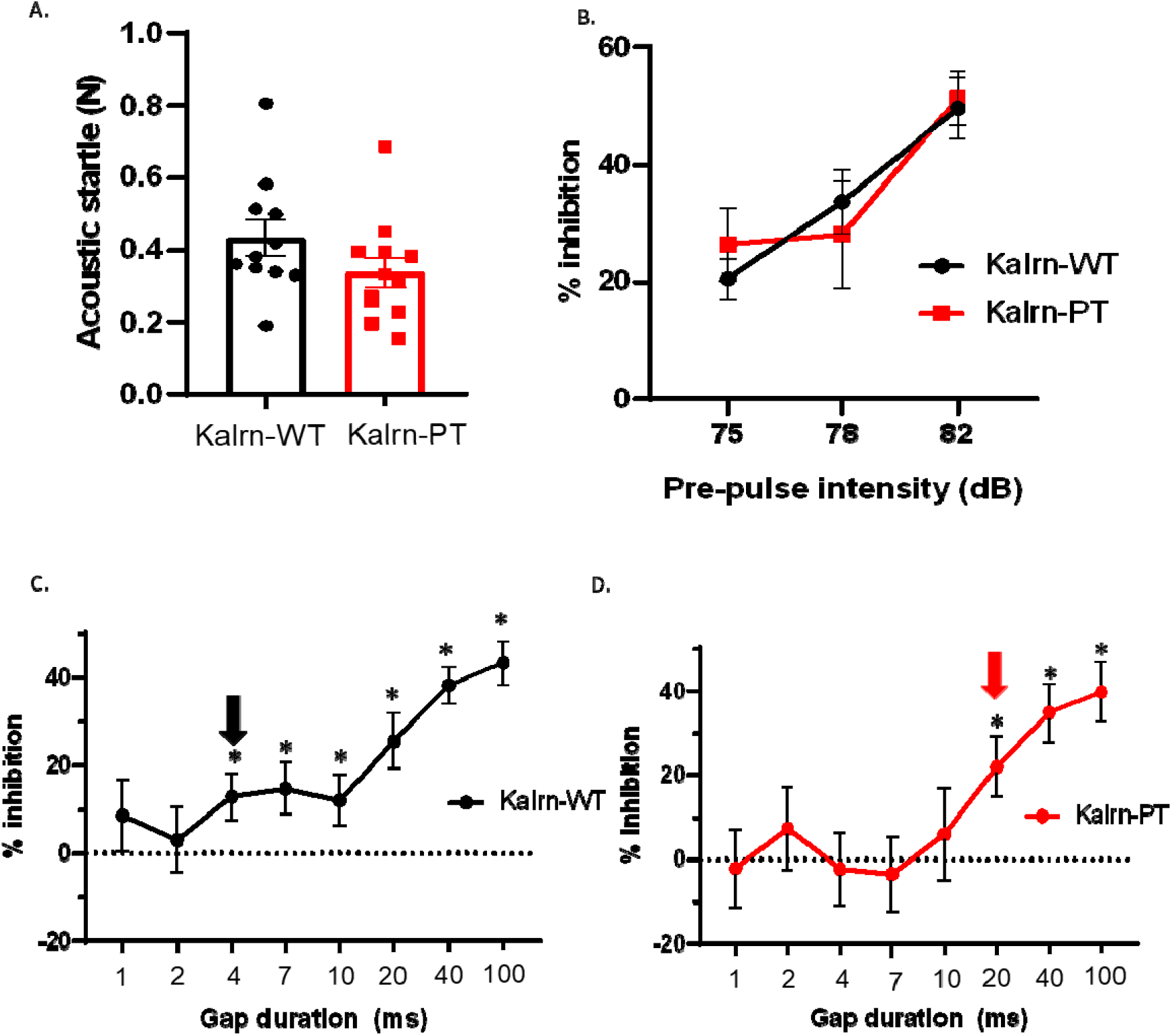
*Kalrn*-PT demonstrate impaired detection of short gap duration without loss of noise-PPI. (A) Baseline acoustic startle response was measured for each animal on day 1 of testing. No difference was observed between genotypes in baseline startle (p=0.15). (B) Both *Kalrn*-WT and *Kalrn*-PT mice demonstrate increasing inhibition to increasing pre-pulse intensity (p<0.0001). Noise-PPI was not different between genotypes (p=0.91). (C) Gap-PPI responses in *Kalrn*-WT mice show increasing inhibition as a function of increased gap duration. Gap detection threshold is defined as the first of at least 2 consecutive points which significantly differ from baseline. Arrow at 4ms indicates gap detection threshold in *Kalrn*-WT mice (p=0.031 at 4ms). (D) Gap-PPI responses in *Kalrn*-PT mice show increasing inhibition as a function of increased gap duration, but detection threshold as defined above is shifted to 20ms (p=0.011 at 20ms). All data shown are ± SEM. *p<.05

### *Kalrn-PT* exerts effects across development

Despite onset of clinical symptoms being delayed until late adolescence/early adulthood, schizophrenia is considered a neurodevelopmental disease. The neurodevelopmental hypothesis of schizophrenia has received support from recent research that implicates shared genetic risk with other syndromes typically considered “neurodevelopmental disorders” with childhood onset (autism spectrum disorder, intellectual disability), and the high degree of co-morbidity suggests an underlying pathology for schizophrenia that targets early brain development (63). Dendritic arborization is known to occur early in development, with arbors reaching near adult-size by P21 in mice and subsequent changes acting primarily to refine circuitry (64, 65). Thus, to determine if the dendritic impairment observed in the 12-week cohort arose early in development, we generated a cohort of 4-week ± 3 days mice. Six neurons/animal were imaged and manually reconstructed in NeuroLucida, and a total of 5 animals/sex/genotype was included. Similar to the 12 week cohort, there was no effect of sex on any of the dendritic measures and thus the sexes were combined to provide 10 animals/genotype in the final analyses. We analyzed the age effect within genotypes and found a significant genotype*age interaction for both length and intersections. Specifically, basilar dendritic length (Fig 8C) and complexity (Fig 8A) remained stable across development within *Kalrn-WT* mice. Conversely, there is a regression in basilar arbor length (Fig 8C) and complexity (Fig 6A) within *Kalrn-PT* mice across development. In contrast, apical arbors exhibit modest continued growth across adolescent development (Fig 8B,D), consistent with previous reports of L3 PCs across adolescence in rodents(26). In *Kalrn*-PT mice, however, the arbors remain static in length and complexity across adolescence (Fig 8B,D). Sholl analyses for each parameter measured within genotype across ages are shown in SI appendix (Fig S5).

**Figure 8.**
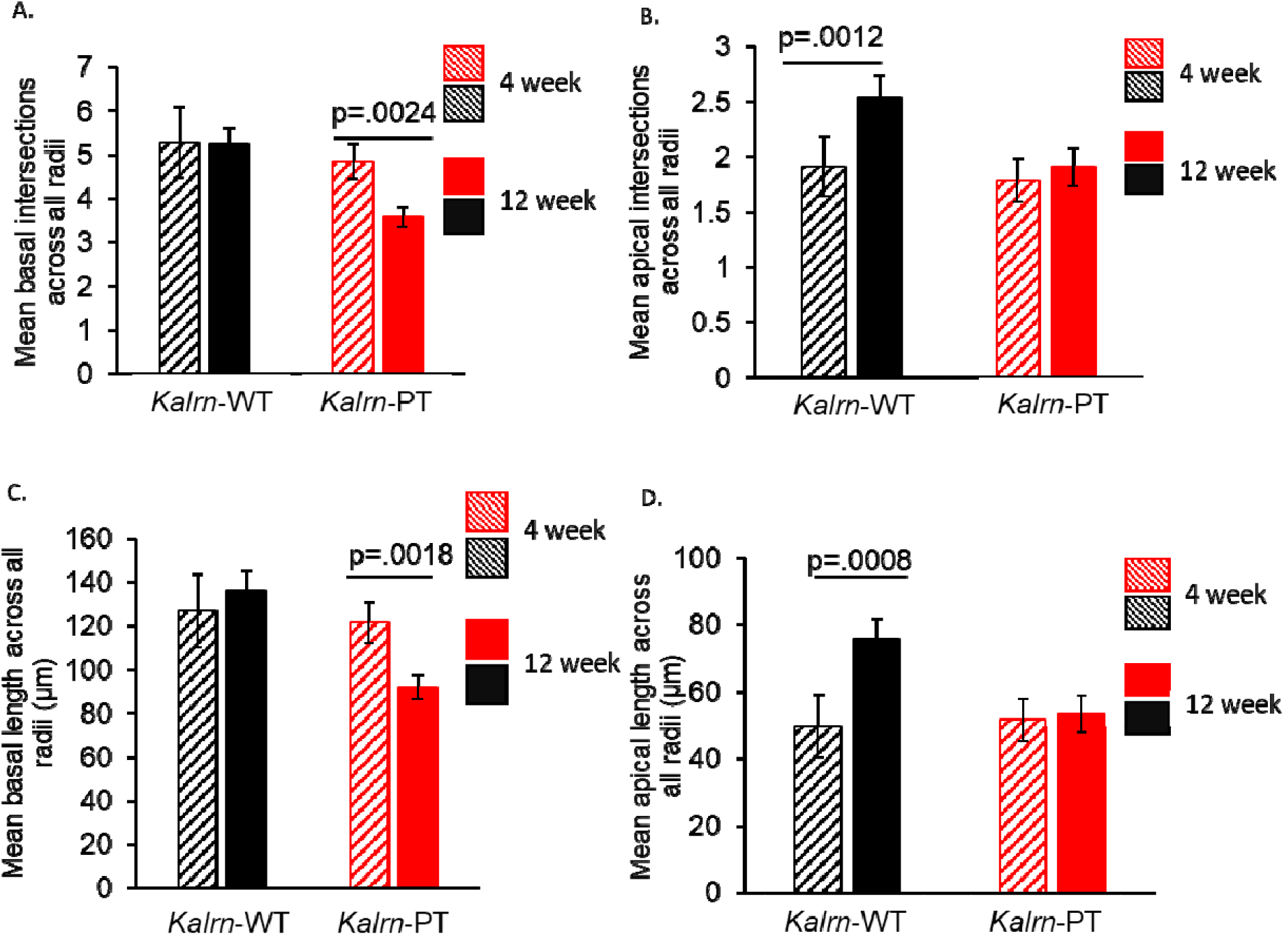
Changes in dendritic architecture across adolescence in *Kalrn*-WT and *Kalrn*-PT mice. (A) Mean basilar intersections across age groups demonstrate a significant regression across adolescence in the *Kalrn*-PT animals, with no change in *Kalrn*-WT animals. (B) Mean apical intersections across age groups shows a significant increase in complexity in *Kalrn*-WT animals, with no change in *Kalrn*-PT animals. (C) Mean basilar length across age groups shows a regression across adolescence in *Kalrn*-PT, but not *Kalrn*-WT, animals. (D) Mean apical length increases across adolescence in *Kalrn*-WT animals, with no change in *Kalrn*-PT animals. All datasets shown are from the previously defined cohorts, N=10 animals/genotype. Data are displayed ± SEM. p-values for individual arbor analyses are displayed above the graphs. Total arbor statistics shown are *p<0.1

Fig 9 shows the results of the Sholl analysis from the 4-week cohort. Unlike in the 12-week cohort, there were no observed differences by genotype across dendritic length or branching at 4-weeks (p-values ranging from p=0.304-0.904 across parameters). Representative reconstructions are shown in Fig 9A,B.

**Figure 9.**
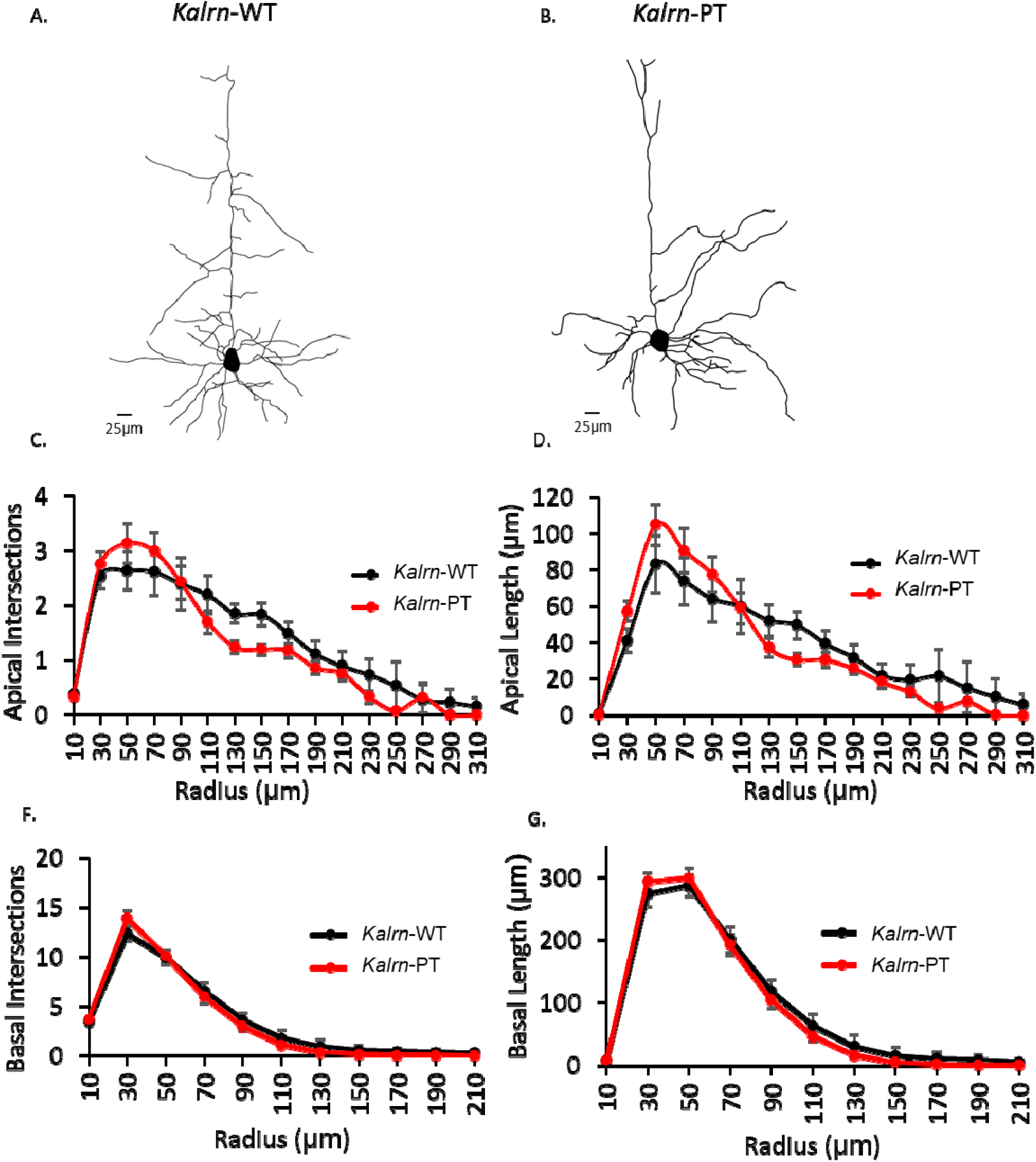
Dendritic arbors form normally in pre-adolescent *Kalrn*-PT mice. Dendritic reconstructions from Golgi stained PCs in A1 of 4-week old WT (A) and *Kalrn-PT* (B) mice demonstrate no change across dendritic complexity or length in (C,D) apical or (E,F) basal arbors. (N=10 mice per genotype, 5 neurons per animal) Results shown are from repeated measures ANOVA with main effect of genotype. Data shown are average values across animals ± SEM.

## DISCUSSION

We hypothesized that the *Kalrn-*PT mutation expressed in KAL9 (KAL9-PT) would increase baseline RhoA activity in neurons, specifically within the subcellular distribution consistent with the protein’s normal expression. We further hypothesized that this activity would be downstream of NGR1 signaling and have functional effects on dendritic morphology. Using genetically encoded RhoA sensors, we find increased baseline RhoA activity in the proximal dendritic shaft in cortical neurons overexpressing KAL9*-* PT. We further found that genetic knockdown of KAL9 mitigates the structural impairment seen in dendrites with NGR1 overexpression. We demonstrated for the first time that OMGp acts via NGR1 in PCs to restrict dendritic length and complexity, an effect which requires KAL9 and is enhanced by KAL9-PT. In a genetic mouse model of the *Kalrn-*PT mutation, we found impairments in dendritic morphology of L3 pyramidal neurons in auditory cortex that resulted from dendritic arbor regression across adolescence. These dendritic structural impairments lead to reduction in spine density per unit of gray matter volume in the absence of changes in spine density per unit of dendrite length. Finally, we show that early adult *Kalrn*-PT mice demonstrate an impaired auditory gap detection threshold.

### KAL9 as a spatially restricted RhoGEF

*Kalrn* belongs to a conserved family of proteins that possess GEF domains. The KAL9 isoform, a dual GEF containing 2 active domains, is largely excluded from dendritic spines (30), instead localizing primarily to the dendritic shaft. This distinct subcellular localization suggests it tightly regulates RhoA activity within a specific microdomain within the neuron. KAL9 likely exerts its effects primarily through cytoskeletal rearrangements within the dendrites. Our findings in neurons support the notion that KAL9 regulates RhoA activity in dendrites, as we see increased RhoA activity in the proximal shaft, but not soma, of neurons overexpressing KAL9-PT. Correspondingly, the most pronounced disparity in both dendritic length and dendritic intersections between *Kalrn-*WT and *Kalrn*-PT occurs between 30-50µm from the soma, which corresponds to the ROIs with the greatest increase in baseline RhoA activity.

The role of RhoA activation in dendritic biology is complex. Although constitutively active RhoA reduces dendritic morphogenesis in rodent PCs(31, 32), expression of dominant negative RhoA fails to affect dendritic outgrowth during development(33). These finding suggest RhoA activity may be significantly restricted during dendritic development and modulation of specific effectors is required to block its activity. Previous work in cerebellar granule neurons identified KAL9 as a critical component of the NGR1/p75 signaling pathway, with KAL9 being responsible for activation of RhoA in neurites(34). However, given NGR1’s numerous signaling mediators in different cell types in the brain (68), we sought to determine if KAL9 regulated NGR1 specifically in pyramidal neurons. To test this, we performed a series of experiments involving both overexpression as well as genetic knockdown approaches. The effect of NGR1 overexpression on dendritic morphology in CA1 hippocampal pyramidal neurons has been extensively characterized (45), allowing us to utilize this system to test if KAL9 is necessary to mediate NGR1’s effects. We show for the first time that KAL9 is necessary for the effect of NGR1 overexpression on dendritic architecture in CA1 pyramidal neurons, as genetic knockdown of KAL9 rescues the structural impairments induced by NGR1 overexpression. Next, we demonstrate for the first time that the MAI OMGp acts as a ligand for NGR1 to restrict dendritic length and complexity. OMGp’s inhibition of dendritic growth is reversed by genetic ablation of KAL9, suggesting KAL9 is required for OMGp’s effects on dendritic architecture. Interestingly, KAL9-PT enhances OMGp’s inhibition of dendritic growth, suggesting increased RhoA activation may be a downstream mechanism by which OMGp exerts its effects. While KAL9 likely signals through numerous pathways(69), it is unlikely that an orthogonal pathway is responsible for these findings as genetic ablation of KAL9 alone does not cause a significant change in dendritic architecture compared to control.

### Dual GEFs and neuropsychiatric disease

Rho GTPases have emerged as key regulators of dendritic morphogenesis and, not surprisingly, are increasingly found to be associated with neuropsychiatric diseases whose pathology includes abnormal dendritic architecture. For example, a large proportion of genes implicated in intellectual disability encode small GTPases (68, 69). More recent evidence further suggests that disruptions in regulators of small GTPase activity, namely dual GEFs, are associated with human neurodevelopmental diseases. A hotspot of *de novo* mutations in Trio, a dual-GEF paralogue of KAL9, was recently described in autism-spectrum disorder (70). An exome-wide significant enrichment in Trio loss-of-function mutations has also more recently been described in schizophrenia (https://schema.broadinstitute.org/gene/ENSG00000038382). *KALRN* itself has been implicated in multiple neuropsychiatric diseases, perhaps most notably schizophrenia (71). The *KALRN*-PT mutation was initially discovered in a re-sequencing analysis of a schizophrenia cohort, conferring an OR>2 (29).

More recently published evidence found a transcriptome-wide increase in exon skipping in *KALRN* transcripts in schizophrenia (51). The exon skipped at a greater rate in schizophrenia was exon 36, present only in the dual GEF isoforms, KAL9 and KAL12. The increasing body of evidence implicating dual GEFs in human neurodevelopmental disease, in particular schizophrenia, underscores the potential importance of tightly regulating small GTPase activity with precise spatial and temporal resolution.

### Dendrites in disease and development

A consistent feature across the above-mentioned neurodevelopmental disorders, most notably schizophrenia, is impaired dendritic architecture. Specifically, reductions in both dendritic spine number and density as well as dendritic length are among the most reproducible findings in postmortem studies of schizophrenia (10). Since dendrites are a principal component of cortical gray matter, it is possible that their reduction in SZ is a contributing factor to the observed reductions in gray matter volumes in disease. Importantly, baseline gray matter volumes are not reduced in clinical high risk groups between those that will later go on to develop SZ and those that will remit (72-74). Instead, the reductions emerge during adolescence and young adulthood concurrent with the transition to symptomatic illness.

Thus, understanding molecular mechanisms leading to impairments of dendritic growth, development, and maintenance in SZ requires developmental context. While overexpression and knockdown models are powerful tools in allowing us to understand the basic biology of molecules with respect to active domains, signaling pathways, etc, they fail to recapitulate normal developmental patterning and physiologic levels of expression. CRISPR/Cas9 affords the ability to selectively edit the genome with high precision, including inserting single base pair mutations at a specific locus (75, 76). Thus, we employed this modality to generate a mouse line containing the *Kalrn*-PT missense mutation at the endogenous locus to better study the effects of modest gain-of-function RhoA activity in *Kalrn* across development.

In mice the final form of the dendritic tree is largely established by the first 2 weeks of postnatal life, well prior to adolescence (17). Afterwards, large-scale structures are remarkably stable, although small degrees of growth have been observed in L2/3 pyramidal neurons using long-term imaging approaches (18). Results from genetic knockdown studies *in vivo* suggest that mature cortical neurons retain the ability for growth but typically remain stable in size due to competing inhibitory pathways (17). Our data support this notion, as we show that apical dendrites are capable of continued growth across adolescence in WT mice, albeit only a modest increase.

Our data from the *Kalrn*-PT mice suggest that emergence of a structural phenotype results not from failure of normal dendritic growth early in development but, rather, regression of the existing dendrites during adolescence. Specifically, the dendritic arbors in the *Kalrn*-PT mice appear to reach a normal final form in pre-adolescence, indicating that they undergo the rapid growth phase early in development without interference. However, structural deficits emerge across adolescence and are evident by early adulthood, suggesting a pathological disruption of normal developmental processes. Our data show that apical arbors fail to achieve modest continued growth across adolescence. Since they achieve a normal pre-adolescent form it is unlikely that this failure to grow is a deficit in a growth pathway but, rather, is more likely attributable to increased activity in a regression pathway. This is supported by our observation that the basilar arbors regress across adolescence in *Kalrn*-PT mice whereas wildtype basilar arbors remain stable, again suggesting that *Kalrn*-PT confers a pathological increase in dendritic regression emergent across adolescence.

Previous data suggest a role for MAIs in regulating cytoskeletal dynamics and neurite length (46, 77-79). Thus, an important consideration is whether the adolescent onset of dendritic regression seen in our *Kalrn*-PT mice reflects enhanced OMGp signaling as seen in our *in vitro* studies. OMGp is expressed both on the membranes of oligodendrocytes as well as by mature neurons(41, 47). In mouse, OMGp increases from P0 onwards, reaching maximum levels in adulthood (41). This developmental patterning of expression poises OMGp to play a key role across adolescence and into adulthood, likely functioning in normal dendritic growth restriction and stabilization. The synergistic interaction of increasing expression of OMGp across normal adolescence with a genetic vulnerability (in this case, *Kalrn*-PT) that enhances RhoA signaling downstream of NGR1, would then result in dendritic regression emerging by early adulthood.

### Dendrites, auditory processing integrity, and schizophrenia

Dendritic length and branching determine a PCs receptive field(11, 12), help to segment computational compartments(13), and contribute substantially to how the received signals are integrated and transmitted to the cell body(14, 15). Dendrites in early adult *Kalrn*-PT animals demonstrate less length and complexity than their wildtype counterparts. Notably, this pathology also results in reduced dendritic spine tissue density due to the dendritic arbor changes themselves, as there was no accompanying change in spine density per unit length of dendrite. Postmortem findings in schizophrenia similarly show a reduction in dendritic length and in dendritic spine tissue density(10). Dendritic spines receive the majority of glutamatergic synapses in the cortex. In the auditory cortex, most of the spines receive narrowly tuned frequency inputs and, as such, serve to spatially and biochemically segregate frequency tuning (80). The reduced numbers of spines would therefore be hypothesized to impair frequency discrimination and processing of auditory information, ultimately producing the auditory sensory processing impairments seen in schizophrenia (5).

One of the most reproducible EEG findings in SZ is decreased amplitude of the mismatch negativity response a change in sound properties, such as silent gaps in noise(6, 81). Gap detection, which in rodents can be indirectly assessed via the inhibition of the acoustic startle response when a prepulse, consisting of a gap in a noise background, is detected, requires integrity of the auditory cortex. In contrast noise-PPI, reflects sensorimotor gating that relies on subcortical circuitry and can still be elicited when the cortex is silenced or lesioned(61, 62). We found that *Kalrn*-PT mice have reduced inhibition specific to short gap duration (4ms in *Kalrn*-WT versus 20ms in *Kalrn*-PT), indicative of an impairment in gap detection threshold. The inability to detect short gap durations suggests a very specific auditory information processing deficit. These findings provide a useful model in which structural impairments within auditory cortical neurons are associated with a specific auditory cortical processing deficit, rather than a global impairment in sensorimotor gating.

### Conceptualizing genetic vulnerabilities across normative development

The data presented within show that *Kalrn*-PT induces adolescent-onset dendritic regression via enhancement of NGR1 signaling to RhoA. These findings contribute to the expanding understanding of how alterations in the dual GEFs Kalirn, and its paralog, Trio, contribute to risk for schizophrenia. Moreover, we provide a model for how a mild genetic vulnerability may interact with the normal developmental timeline such that pathology only emerges in or after adolescence. This interplay between genetic susceptibility and normal adolescent development, both of which possess inherent inter-individuality variability, may predict the heterogeneity seen in phenotypes in human neuropsychiatric disease.

## METHODS

### Dissociated cell culture

Rat cortical neurons were prepared from E18 Long-Evans rat embryos (Charles River) as previously described(82). In 24-well plates, neurons were plated at 1.6×10^5^ cells/well on glass coverslips (acid washed) coated with poly-D-lysine (20 μg/mL) and laminin (3.4 μg/mL). Cortical neurons were maintained in Neurobasal Medium (Invitrogen), supplemented with 2% B27 (Invitrogen), penicillin/streptomycin (100 U/mL and 100 mg/mL, respectively), and 2 mM glutamine with fresh media supplementation every 4 days. Dissociated neurons were transfected at DIV8 using Lipofectamine 2000 (Invitrogen) according to the manufacturer’s suggestions. Wildtype KAL9 (KAL9-WT) or mutant KAL9 (KAL9-PT)(30) was added at 600ngs. Genetically encoded RhoA sensor construct (45) was added at 350ngs and PCS2 filler DNA(45) was added up to 1 µg, along with 2µL Lipofectamine 2000 (Invitrogen) per well. For sensor imaging, neurons were placed in mACSF(45). Imaging was performed at DIV14-18. A total of 3 independent cultures were generated, with 8-10 neurons imaged per condition within each experimental round.

For dendritic arborization experiments *in vitro*, cultures were derived as above. Dissociated neurons were transfected at DIV8 as above with vGFP (250ng/well) and either shNGR1 (10 ng/well)(45), shKAL9 (15 ng/well) (34), KAL9-WT (600 ng/well) or KAL9-PT (600 ng/well). Commercially available recombinant mouse OMGp (R&D Systems) was reconstituted to 100 µg/mL in sterile PBS. On DIV12 cultures were treated with either sterile PBS alone or 200 ng of OMGp per well of a 24-well plate. Cultures were fixed at DIV14 in 4% PFA and mounted using Fluromount-G (Invitrogen). A total of 3 independent cultures were generated per experimental condition, with 8-10 neurons imaged and reconstructed per condition within each experimental round.

### Organotypic Slice Culture

Transverse slices (400 μm) of P6 hippocampus from Long-Evans rats were prepared and cultured essentially as described in (Stoppini et al., 1991). Slices prepared under sterile conditions and were cultured on hydrophilic PTFE inserts (0.4 μm pore size, Millicell) in 6-well dishes containing 0.75 mL of antibiotic-free medium (20% horse serum/MEM) and incubated in 5% CO2 at 37°C. Slice cultures were transfected using a Helios Gene Gun (Biorad) at DIV4. For each bullet, up to three different plasmids comprising a total of 50 μg of DNA (10 μg shKAL9 construct(34), 20µg WTNGR1(45),15 μg vGFP) were coated onto 12.5 mg of 1.6 μm gold particles, as per the manufacturer’s protocol (Biorad). Slices were fixed at DIV11 in 2.5% paraformaldehyde and 4% sucrose and processed for immunofluorescence as previously described(44). In brief, slices were counterstained with Hoechst (1:800) and chicken polyclonal anti-GFP (Aves Labs, 1:800), and the nylon inserts were mounted onto glass slides using Fluormount-G (Southern Biotech). A total of 3 independent cultures were generated, with 5-10 neurons imaged per condition within each experimental round.

### In vitro Imaging

Imaging for slice cultures and sensor studies were performed using a Nikon A1R laser scanning confocal. Sensor imaging studies in dissociated cortical neurons were done as described previously(44). Experimenter was blind to condition throughout all image acquisition, reconstruction, and analysis. Approximately 8 neurons were imaged per condition for each experiment, and each experiment was performed three times. Sholl analysis, as well as measures of average total dendritic length and complexity, were carried out for both apical and basal dendrites using NeuroLucida and NeuroExplorer software. For dissociated culture experiments, images were captured on an upright epifluorescent microscope (Olympus BX53 with UPlanSApo 10x objective, 1.29 µm/pixel) and manually reconstructed using NeuroLucida software. Sholl analyses were conducted in NeuroExplorer. Experimenter was blind to condition throughout all image acquisition, reconstruction, and analysis. 8-10 neurons were imaged and reconstructed per condition within each experimental round, for a total of 24-30 neurons per condition across 3 independent experimental runs.

### CRISPR/Cas9

#### CRISPR Reagent Production

A truncated sgRNA (84) targeting *KALRN* was designed using CRISPR Design Tool (85). A sgRNA specific forward primer and a common overlapping reverse PCR primer (see SI appendix, Table S1) were used to generate a T7 promoter containing sgRNA template as described (86). This DNA template was transcribed *in vitro* using a MEGAshortscript Kit (Ambion). The Cas9 mRNA was prepared using a MEGAshortscript kit (Ambion) as described (87). Following synthesis, the sgRNAs and Cas9 mRNA were purified using the MEGAclear Kit (Ambion), ethanol precipitated, and resuspended in DEPC treated water. A single stranded 130 nucleotide DNA repair template (“repair oligo,” see SI appendix, Table S1) harboring the knockin sequence (C→A) as well as a silent mutation (C→T) introducing a unique Hinf1 restriction site was purchased as PAGE purified Ultramer DNA (Integrated DNA Technologies, Coralville, IA). Final codon change was from CCC (wildtype) to ACT (knockin). The oligo was complementary to the nontarget strand (88).

#### Knockin Mouse Production

sgRNA (50ng/µL), Cas9 mRNA (75ng/µL), and the repair oligo (100ng/µL) were mixed and injected into the cytoplasm of C57BL/6J one-cell mouse embryos as described (87).

#### Mouse Genotype Analysis

Mice were genotyped by PCR amplification (see Si appendix, Table S1 for primer sequences, KALF2 and KALR2) followed by restriction digestion with Hinf1 (FastDigest) to produce PCR amplicons that were either 302bp (wildtype) or 138bp + 164bp (knockin). Undigested PCR amplicons of founder and all F1 offspring were also analyzed by Sanger sequencing.

#### Off target analysis

The top 18 predicted off-target sites (see SI appendix, Table S2) using COD (Cas9 Online Designer) software(89) were analyzed by PCR amplification followed by Sanger sequencing.

### qRT-PCR and Western Blotting

For mRNA collection, frontal pole gray matter was collected in Trizol (Life Technologies) and tissue was manually homogenized by passing through sequentially graded syringe gauges. RNA was isolated using the RNeasy Lipid Tissue kit according to manufacturer’s instructions (Qiagen). RNA was reverse transcribed into cDNA using the Bio-Rad iScript kit, containing a mix of random hexamers and oligo dT primers. Isoform-specific primers were used to detect KAL7, KAL9, and KAL12, as well as the naturally occurring KAL9 splice variant (primers designed spanning the exon 35-37 junction). The housekeeping GAPDH was used for normalization (SI appendix, see Table S4 for primer sequences). Real-time quantitative PCR was performed (95 degrees for 10 min; [95 degrees for 15s, 60 degrees for 1 min] x 40 cycles) on a Step-One Plus (Applied Biosystems) using Sybr Green (Bio-rad) with ROX serving as the internal reference dye. Results were analyzed using the comparative Ct method.

Cell-free protein lysates were obtained from cells using radioimmune precipitation assay buffer (10 mM Tris-HCl, pH 8.0, 1 mM EDTA, pH 8.0, 0.5 mM EGTA, 140 mM NaCl, 1% Triton X-100, 0.1% sodium deoxycholate, 0.1% SDS), and total protein levels were quantified using the Lowry protein assay (Bio-Rad). 15 µg of total protein was loaded per well. Samples were electrophoresed on a NuPage Bis-Tris 4-12% gradient polyacrylamide gel (ThermoFisher Scientific) and transferred to a PVDF membrane (Millipore, Danvers, MA) at 100V for 1h. Membranes were blocked in Licor Blocking buffer (Odyssey, Lincoln, NE) for 1h at room temperature and primary antibody (Kal-spec, Millipore Sigma, Burlington, MA; β-tubulin, Epitomics, Burlingame, CA) was added (1:1000 for Kal-spec, 1:60,000 for β-tubulin) and incubated overnight at 4 degrees Celsius. Membranes were washed 5×5min in TBS containing 0.1% Tween and incubated with fluorescent-conjugated secondary antibodies (1:10,000) for 1h at room temperature. Five more washes were performed and membranes were imaged using the Odyssey Licor System. Densitometric analyses were performed using MCID Core Analysis software (UK).

### Golgi

#### Tissue Generation

Cohorts of mice were sacrificed at age 12 weeks ± 3 days and 4 weeks ± 3 days. These ages correspond in general to young adulthood and pre-adolescence, respectively, in mice (90, 91). Age-matched cohorts that included males and females of the 2 genotypes of interest (wildtype and homozygous *Kalrn-PT*) were processed together throughout. Vaginal cytology was collected on all early adult female mice at the time of sacrifice. Mice were sacrificed by lethal CO_2_ inhalation followed by decapitation and whole brain extraction. Golgi staining was performed using the FD Rapid GolgiStain Kit (FD Neurotechnologies, Inc) according to manufacturer’s instructions.

For the analysis of spine density, we chose to focus on proximally arising secondary apical dendrites for several reasons: 1) We found significant reductions in dendritic length and complexity in apical arbors, demonstrating that *Kalrn-PT* is exerting effects on apical dendrites 2) The focus on secondary apical dendrites allowed us to institute highly consistent criteria for selection. The latter is particularly important given that spine density is known to vary depending on distance from the soma.

#### Auditory Cortex Mapping

Animals were coded so that experimenter remained blind to genotype throughout. Using Slidebook 6.0 (Intelligent Imaging Innovations) software, a 4x montage image of a Golgi-stained coronal section was obtained and aligned with the corresponding Nissl plate containing A1 (which spans approximately 2.18-3.64 mm posterior to Bregma and 4.5mm lateral to the midline) in the mouse brain atlas (Paxinos and Franklin, 2004). An average of 4 sections per animal contained A1. An outline of A1 was drawn at 4x magnification, using the dorsal extent of the posterior commissure as an anatomical fiduciary (SI appendix, Fig S2). Within A1, deep layer 3 was defined as 30-50% of the distance between the pial surface and white matter border (SI appendix, Fig S2) and sampling of pyramidal cells was performed as described below.

### Immunohistochemistry (IHC)

An additional cohort of mice aged 12 weeks ± 3 days was generated. Mice were sacrificed as above, the brains were sagittally hemisectioned, and the left hemisphere was fixed in 4% paraformaldehyde overnight at 4°C. Tissue was then transferred to 18% sucrose solution for sectioning on a cryostat. The cerebellum was removed and the tissue block was mounted and cut rostral to caudal at 40µm thick sections. Every 8^th^ section was pulled for Nissl staining. Nissl sections were used for guidance to determine tissue sections that contained auditory cortex. Detailed methods from our lab for auditory cortex mapping are found in (9). One section containing auditory cortex per animal was used for IHC. In order to visualize dendritic spines, we used two markers in combination: a rabbit polyclonal antibody directed against spinophilin (Millipore AB5669) at a dilution of 1:1500. The second was the f-actin binding mushroom toxin phalloidin (Invitrogen A12380) conjugated to Alexa Fluor® 568. Spinophilin is highly enriched in spine heads (57, 59). Phalloidin binds f-actin which is also highly enriched in dendritic spines (58).

### Behavioral Assays

All behavioral testing was conducted in the Rodent Behavior Analysis Core of the University of Pittsburgh Schools of Health Sciences. A cohort of 12 week ± 3 days animals was generated, comprising 11 *Kalrn-* WT (6F, 5M) and 12 *Kalrn*-PT (6F, 6M) animals. Mice were balanced across testing days for genotype, and prior to testing mice were acclimated to the testing equipment. Mice underwent acoustic startle and noise-PPI testing on Day 1, followed by Gap-PPI testing on Day 2.

#### Acoustic Startle Reflex and Noise-PPI

The acoustic startle reflex (ASR) is an involuntary response that occurs after exposure to a startle-eliciting acoustic stimulus. To assess this reflex, mice were placed on a force-sensing piezoelectric transducer enclosed within a sound attenuating chamber (14 x 19.5 x 10.875 in, −35 +/-2 dB attenuation) (Kinder Scientific, Poway, CA). The platforms were calibrated to 1 +/-0.05 N prior to each session. Acoustic stimuli (full range white noise) were delivered via a built-in speaker centered above the animal enclosure. Noise stimuli were calibrated using a digital sound pressure level meter (model 33-2055, accuracy +/-2 dB at 114 dB, range 50-126 dB, RadioShack, Fort Worth, TX). Animals were able to turn freely during the session, but they were prevented from rearing by the height of the animal enclosure. The ASR was assessed by measuring the reflexive response of mice presented with a 115 dB noise, startle eliciting stimulus, presented against a 65-dB white noise background. The mouse’s startle response, which was measured as the displacement on the force sensing platform, was sampled in 1-ms bins for 70 ms starting at the onset of the startle-eliciting stimulus. The maximum startle response during the 70-ms recording window was then determined automatically by the Startle Monitor software (Kinder Scientific). PPI is the attenuation of the ASR that occurs when a nonstartling stimulus precedes the startle eliciting stimulus(93). To assess noise-PPI, we measured startle responses to the 115-dB startle eliciting stimulus when preceded by 75, 78, and 82 dB noises for a 40ms duration, presented 100 ms prior to the startle eliciting stimulus.

#### Gap-PPI

To assess detection of silent gaps embedded in background noise, we used a modified PPI of the acoustic startle reflex paradigm (Gap-PPI)(94, 95). In this paradigm, the prepulse stimulus is replaced with a silent gap. If the animal detects the gap, the startle response is inhibited, and longer duration gaps are more salient and elicit greater inhibition of the startle response. Startle responses to silent gaps (1, 2, 4, 7, 10, 20, 40, or 100 ms in duration) embedded in a 70-dB white noise background were determined.

### Statistical Analysis

For the RhoA sensor data and the Sholl analyses from slice culture, a repeated measures 2-way ANOVA with pairwise comparison by Bonferroni posthoc was performed in GraphPad Prism 8.0. For the *in vitro* dissociated culture data, a repeated measures one-way ANOVA was performed with Sidak multiple comparisons testing in GraphPad Prism 8.0. Statistical analyses were restricted to radii in which ≥50% of the neurons had non-zero values.

For basilar, apical and total arbor from the *in vivo* Sholl analyses, we aggregated the neurons within the same radius for each animal. Then we plotted the average length or intersection count, averaged across all radii between 30µm and 190µm for each week-by-genotype combination. We also plotted the trajectory of length or intersection count over radius for each genotype group, separated by week.

A linear-mixed effects model (96) was fitted for each length or intersection count, where radius (treated categorical), sex, genotype, week, and genotype-by-week interaction were treated as the fixed effects, and mouse was treated as the random effect. For the 12-week *in vivo* data, instead of the 2-category sex, a 4-category “sex” including male and three vaginal cytology for the female cohort was included as a covariate in the final model (Si appendix, Table S3). We tested the genotype effect by comparing *Kalrn*-PT vs *Kalrn*-WT within each age group, as well as the age effect by comparing week 12 vs week 4 within each genotype group. The analysis was performed with the PROC MIXED procedure in SAS 9.4.

For the cortical volume estimates, a 2-way ANOVA was performed in IBM SPSS Statistics. Spine number from IHC was analyzed by 1-way ANOVA in GraphPad Prism.

For Gap-PPI analyses, percent inhibition of startle response was calculated for each gap duration as 100*[(mean startle-only trial response – mean gap trial response)/mean startle-only trial response]. For the noise-PPI experiments, percent inhibition of startle response was calculated for each noise prepulse stimulus using a similar equation, 100*[(mean startle-only response – mean noise pre-pulse trial response)/ mean startle-only trial response]. Effects of genotype on both gap- and noise-PPI were tested using repeated-measures ANOVA in SPSS.

Group-level gap detection thresholds (lowest gap duration to elicit significant PPI) were determined by comparing the mean percent inhibition for each gap duration bin against zero using a one-sample t-test in GraphPad Prism. The threshold was defined as the first of at least two consecutive gap duration bins for which the percent inhibition was significantly different from zero.

Please see SI material for expanded methods

## ACKNOWLEDGEMENTS

MH071533 (RAS), NARSAD Distinguished Investigator Grant from the Brain &Behavior Research Foundation (RAS), MH118513-01 (MJG), MH071316 (PP), MH097216 (PP), AA020889 (GEH), R56AG058593 (ZPW), NARSAD Young Investigator Award (ZPW).

