## Supplementary material for "A Kalirin Missense Mutation Enhances Dendritic RhoA Signaling and Leads to Regression of Cortical Dendritic Arbors Across Development": SI appendix

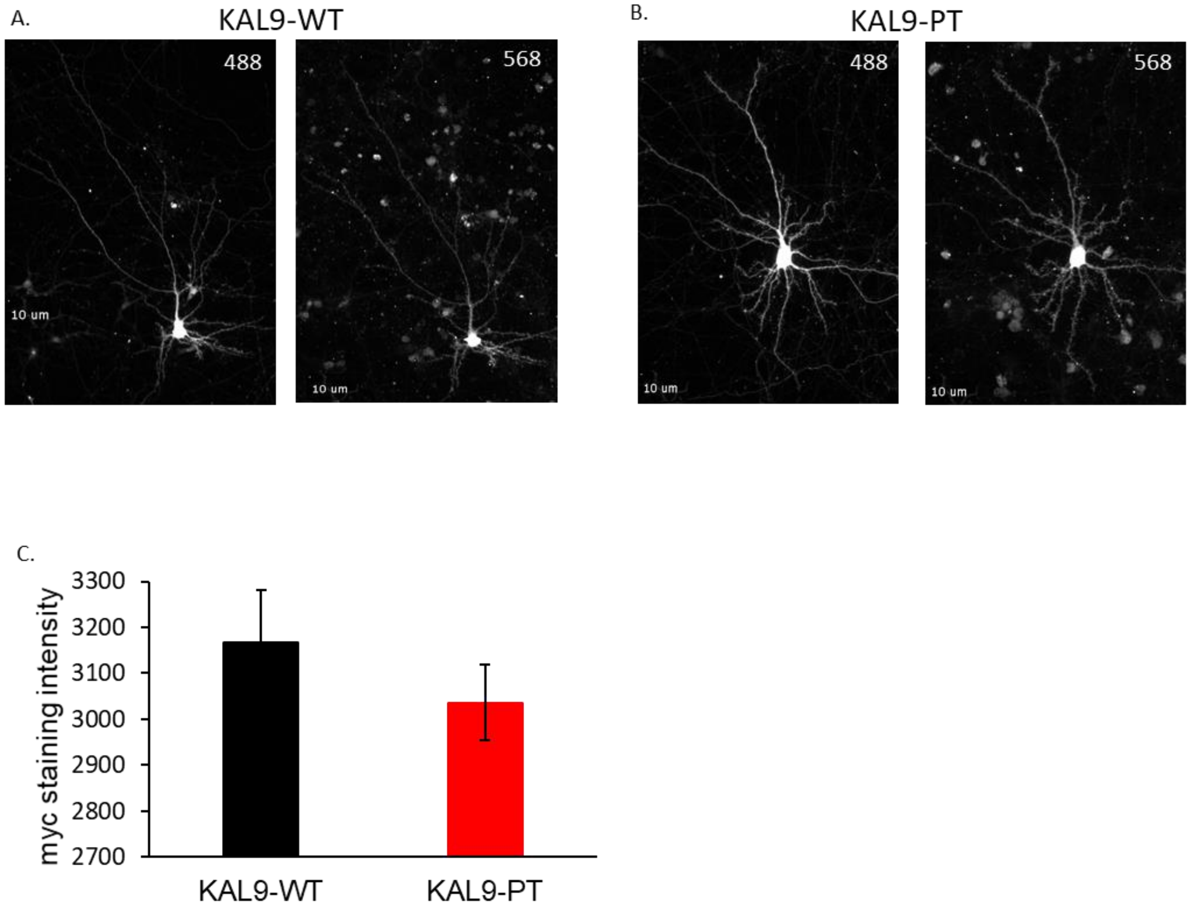


**Figure S1. Overexpression of KAL9-WT and KAL9-PT in dissociated cortical neurons.** Representative images of neurons transfected at DIV8 with vGFP and either (A) KAL9-WT-myc and (B) KAL9-PT-myc. Neurons were fixed and stained for c-myc at DIV14. GFP images were acquired in the 488 channel, an anti-myc antibody was used with a 568-conjugated secondary antibody for myc detection in the 568 channel. Images were acquired at fixed exposure times (25ms in 488, 12ms in 568) at 10x. Ridler-Calvard automatic segmentation masking was used to generate masks in both the 488 and 568 channels. Mask intensities were extracted and background subtracted. (C) Quantitative data shows no significant difference in expression levels between genotypes (KAL9-WT = 3167.28 ± 322.23, KAL9-PT = 3035.99 ± 231.75; p=0.257). N=75 KAL9-WT and N=84 KAL9-PT neurons distributed across 2 independent experiments.

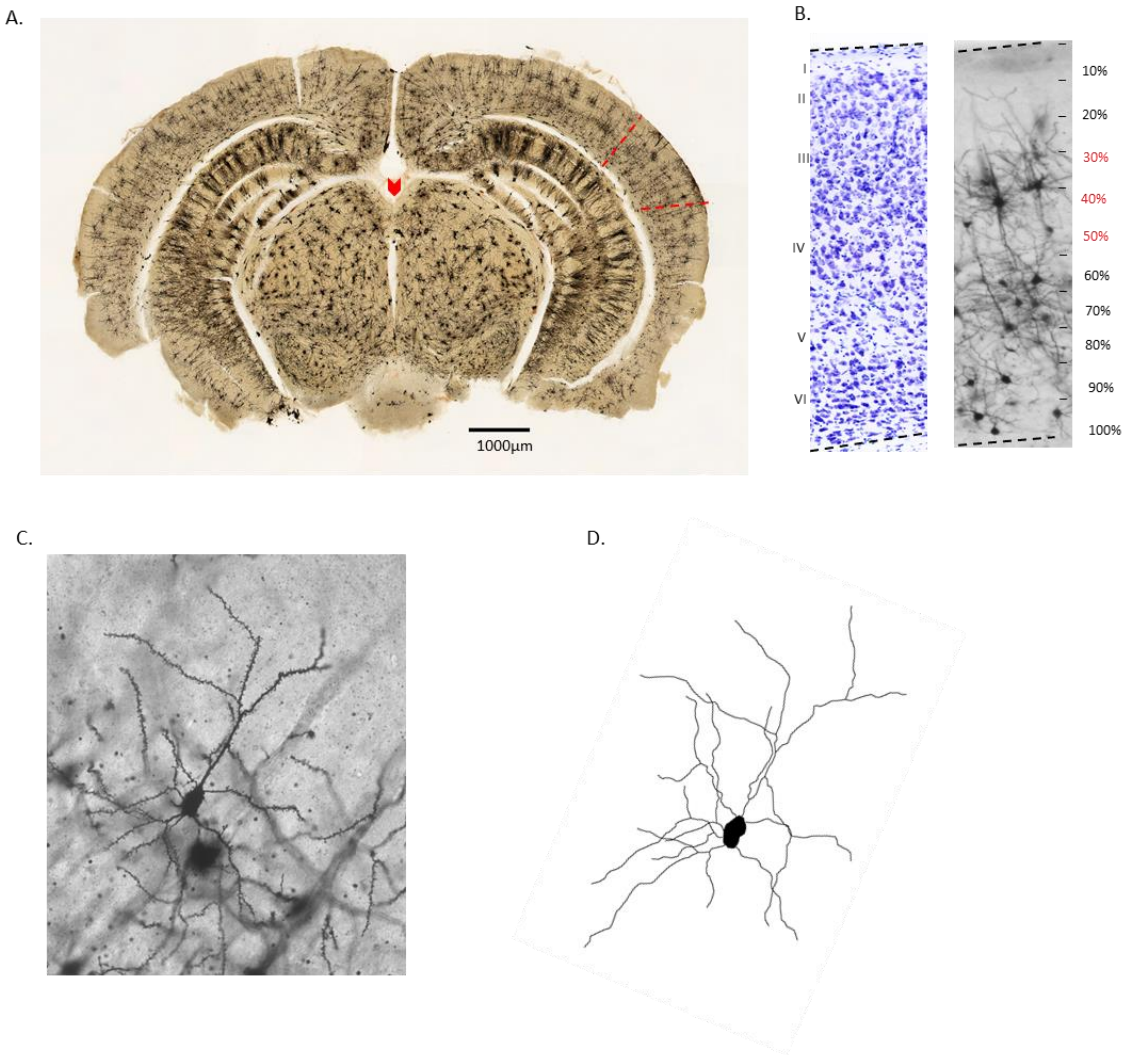


**Figure S2. Identification of Layer 3 Auditory Cortex and Neuronal Reconstructions.** (A) Whole section montage image at 4x magnification of a Golgi-stained section containing A1. Dorsal extent of posterior commissure (indicated by red arrowhead) was used as a gross anatomical landmark, and a horizontal line was drawn to the pia. Using the mouse atlas for reference (plate 55, Bregma -2.92), A1 was contained within the region spanning 1mm superior and 250 μm inferior to this landmark (indicated by red dashed lines). (B) L3 was defined as 30-50% of the distance between the pial surface and white matter border, as determined from alignment with a Nissl-stained section generated for reference. (C) Representative image of a Golgi stained PC at 40x magnification. Images were collected at 1μm step size throughout the image depth. Image shown is a flattening of 10 planes. (D) Neurolucida drawing of the neuron shown in Panel C.

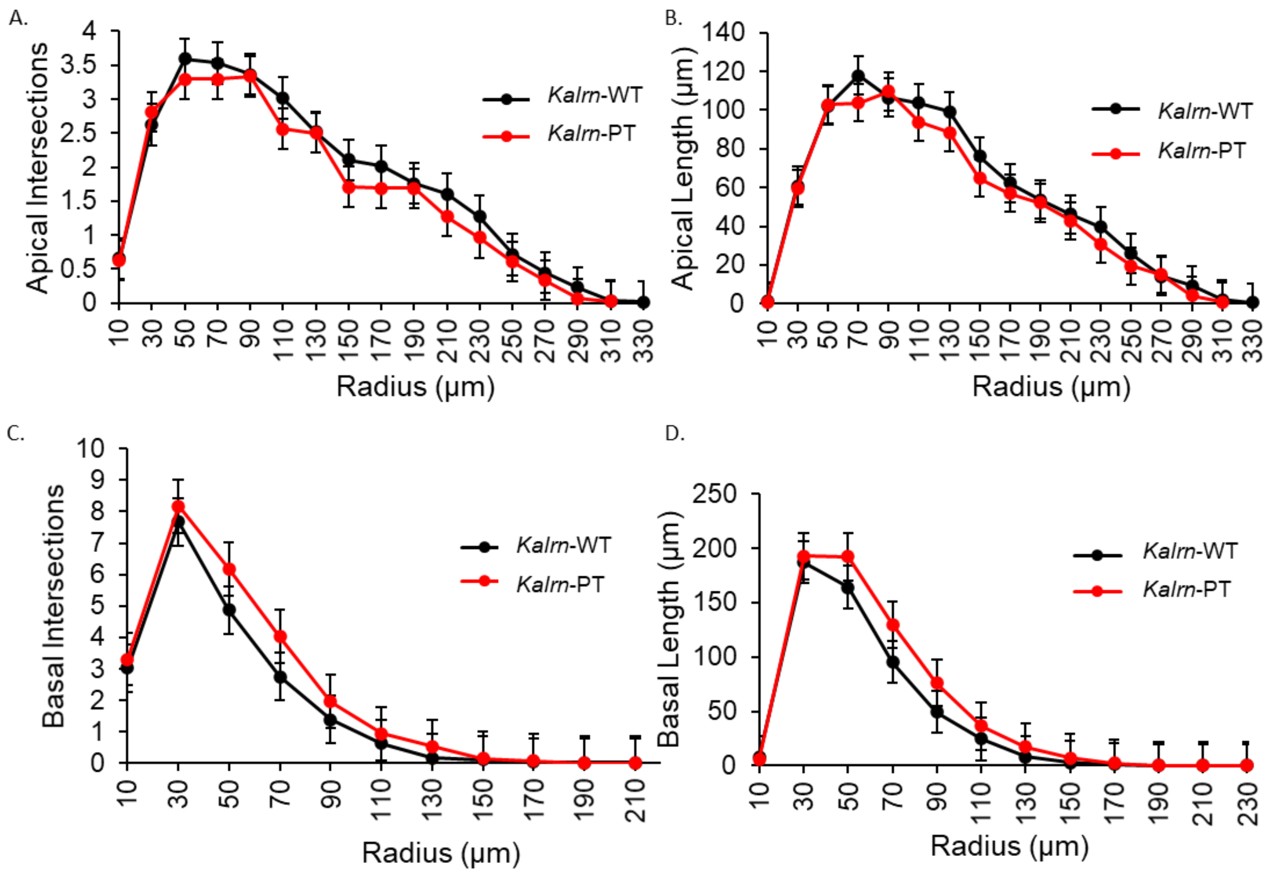


**Figure S3. Sholl analysis of Layer 5 PCs in A1 from *Kalrn*-WT and *Kalrn*-PT mice.** Reconstructions and Sholl analysis of Golgi stained Layer 5 PCs in A1 of 12-week old *Kalrn*-WT and *Kalrn-PT* mice demonstrate no change in dendritic complexity and length in (A,B) apical arbors or (C,D) basilar arbors. (N=10 mice per genotype, 6 neurons per animal) Results shown are from repeated measures ANOVA with main effect of genotype. Data shown are average values across animals ± SEM.

A.

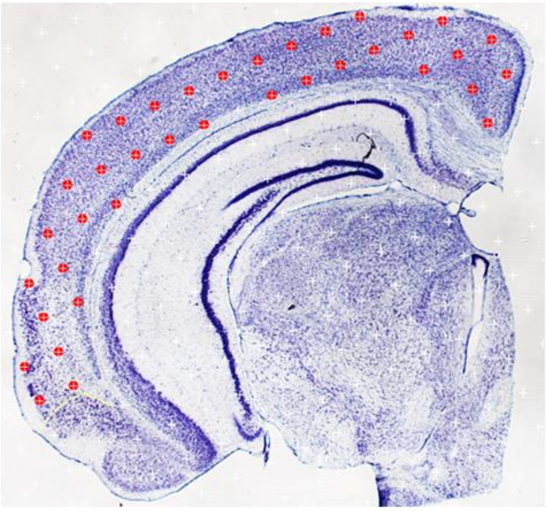

B.

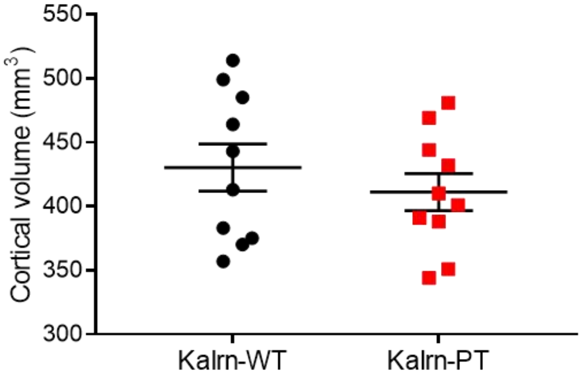


**Figure S4. Determination of cortical gray matter volumes in *Kalrn*-WT and *Kalrn*-PT mice.** (A) Representative image of a Nissl-stained section with overlaid point grid for Cavalieri estimator method. Red points indicate those falling within the defined neocortical region. (B) A small but non-significant trend (p=.24) towards reduced neocortical volume was seen in *Kalrn-PT* mice compared to *Kalrn*-WT (430 $mm^3$ (WT) v. 411 $mm^3$ (PT); N=10 animals/genotype). All data shown are ± SEM.

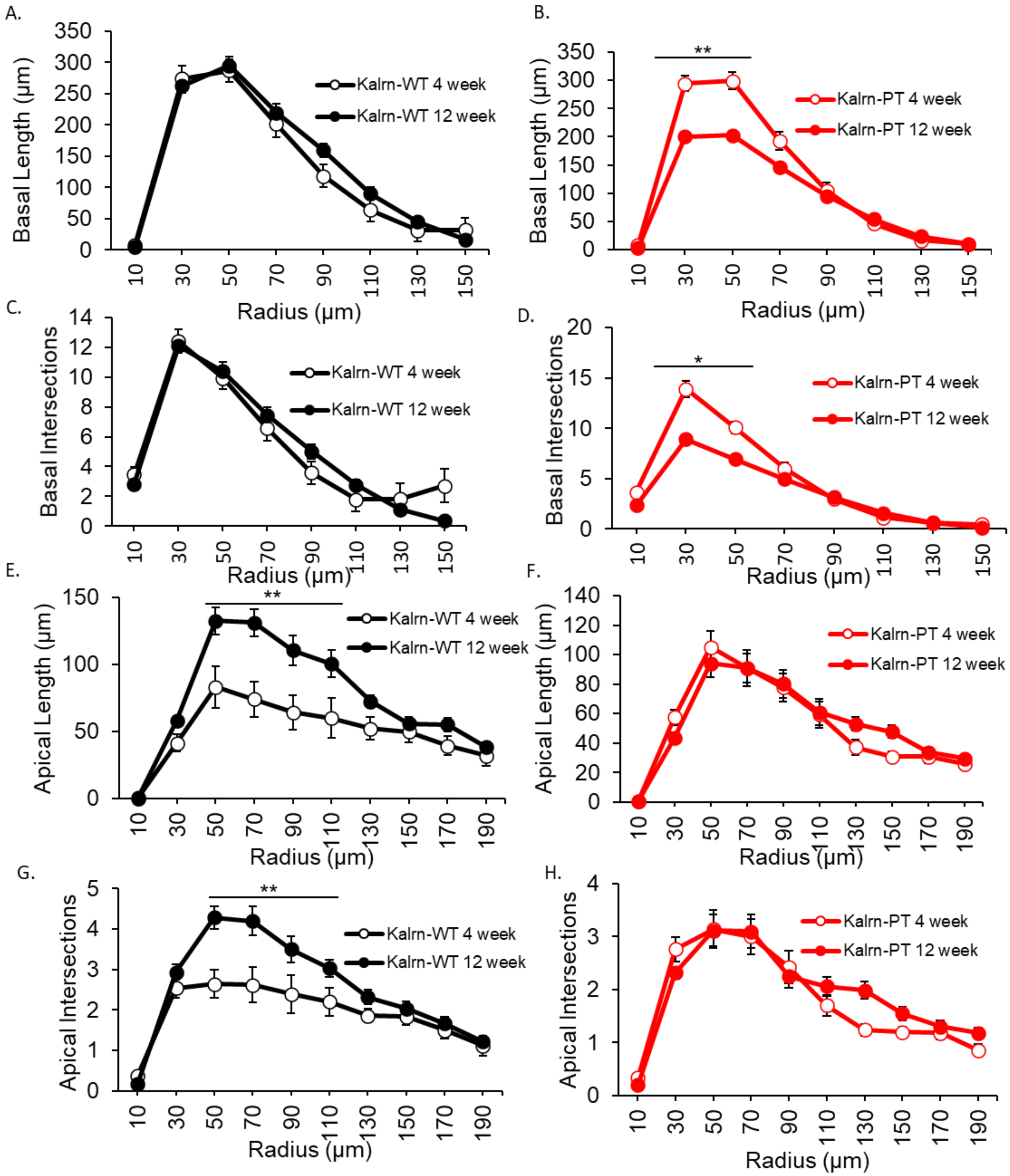


**Figure S5. Sholl analyses across age groups in *Kalrn*-WT and *Kalrn*-PT animals.** (A-D) Basilar arbors were analyzed for both length and complexity across age groups in *Kalrn*-WT and *Kalrn*-PT animals, demonstrating a significant regression in proximal regions between 4 and 12 weeks in *Kalrn*-PT animals. (E-H) Apical arbors were analyzed for both length and complexity across age groups in both genotypes. A significant expansion in proximal regions was noted in *Kalrn*-WT animals. All datasets shown are from the previously defined cohorts, N=10 animals/genotype. Data are displayed ± SEM. *p<.05, **p<.01

**Table S1.** PCR primers and DNA repair templates.
All sequences are written in the 5' to 3' direction.
Sequences in **bold** are T7 promoter. Underlined sequence is the sgRNA target sequence.
Thr knockin codon sequence is marked in red text in repair oligo. (WT codon is CCC)

| PRIMER | SEQUENCE |
|---|---|
| sgRNA94 F primer | **GAAATTAATACGACTCACTATAGG**TTGTGGAACCCTCGGGGGCTGTTTTAGAGCTAGAAATAGC |
| sgRNA common reverse | AAAAGCACCGACTCGGTGCCACTTTTTCAAGTTGATAACGGACTAGCCTTATTTTAACTTGCTATTTCTAGCTCTAAAAC |
| Repair oligo | CACTACAGTCACCCATTGAGTATCAGCGGAAGGAGAGAAGCACAGCTGTGATCCGGTCCCAGACTCCGAGGGTTCCACAAGCCAGCCCCAGGCCCTACTCTTCTGGCCCGGTGGGCTCAGAGAAG |
| KAL F2 | AACTTTGGTACCTAGCCGGC |
| KAL R2 | AACTTGTCCCCAGGCTGATG |

**Table S2.** Off target site analysis.

| KAL off-targets | | TTGTGGAACCCTCGGGGGCTNGG | activity=1 when PAM=NGG | |
|---|---|---|---|---|
| | mouse chromosome | alignment(dots are matched bp) | off-target score(activity compared to the perfect target) | Primers |
| sgRNA94 | 16 | .................G.. | 1 | N/A |
| KAL.OTR1 | 1 | ....A..........A.. | 0.45486 | F: TGCTCAAGTCTCTCCTGCAA<br>R: TGCTCTGGCAGTCCACTAAG |
| KAL.OTR2 | 2 | ..T....A.........G.. | 0.3980025 | F: TCAGGACCTCTGGAAGAGCA<br>R: AGGGCTGACATAGGTCTGGT |
| KAL.OTR3 | 3 | ....A..........T.. | 0.363888 | F: CAATCCAGTCTCAGCGGTCA<br>R: TCTTCGTGGGTACAAACTGCT |
| KAL.OTR4 | 8 | ..C.T...........C.. | 0.318402 | F: CGACTTCATCAGCACTGGGT<br>R: TGCTTCCATAGTTGCAGCGA |
| KAL.OTR5 | 6 | ...CA...........G.. | 0.318402 | F: CCACCGGCCTCTAACTTGTT<br>R: TCAGTGGTCGACTTCCTTCC |
| KAL.OTR6 | 7 | .............G.. | 0.2547216 | F: GCACTGTGTGGTCACTGACT<br>R: GCCTTGCCTGCCTCTCTTTA |
| KAL.OTR7 | 12 | ...A..........G.. | 0.2547216 | F: TCTTTGGCTGCATGGAACCTCT<br>R: AGCTCAGTGTTGGCTCAGTC |
| KAL.OTR8 | 6 | ..A...........G.. | 0.2547216 | F: TCCCGAGGTCACTCACATCT<br>R: ACCAGGGGAGTGTGAGTGAT |
| KAL.OTR9 | 17 | .............G.. | 0.2547216 | F: GTCAGCCGAAACAAGCAAGG<br>R: CACCAACACCTTGCAAGTCG |
| KAL.OTR10 | 11 | ..G...........T.. | 0.2547216 | F: AAGCCGCTGAGCCGTATTTA<br>R: TTGTGAGGATTGGCCCTGTC |
| KAL.OTR11 | 18 | ......A...G.......G.. | 0.2527 | F: CAGCTCATCACAGGCAAGGA<br>R: TCAACATTCAGGGCCCCATC |
| KAL.OTR12 | 12 | ......A...G.......G.. | 0.2527 | F: CAGCTCATCACAGGCAAGGA<br>R: TCAACATTCAGGGCCCCATC |
| KAL.OTR13 | 9 | ......A...G.......G.. | 0.2527 | F: TTGACCTTGTCCTGGACACG<br>R: TCAACATTCAGGGCCCCATC |
| KAL.OTR14 | 15 | ......A...G.......G.. | 0.2527 | F: TCAACATTCAGGGCCCCATC<br>R: TTTTCTCCCACAGGCTGACC |
| KAL.OTR15 | 15 | ......A...G.......G.. | 0.2527 | F: GATCAACATTCAGGGCCCCA<br>R: AAAATGCAGTGGGAGGGTGT |
| KAL.OTR16 | 15 | ......A...G.......G.. | 0.2527 | F: GAGCTTTTCTCCCACAGGCT<br>R: TCAACATTCAGGGCCCCATC |
| KAL.OTR17 | 13 | ......A...G.......G.. | 0.2527 | F: TCAACATTCAGGGCCCCATC<br>R: CAGCTCATCACAGGCAAGGA |
| KAL.OTR18 | 10 | ......G...T.......T.. | 0.2527 | F: TTGCCAGACAACCTCAGTCC<br>R: TCTCGATGGGCGTCTCTTTG |

| Mouse ID | Sex | Genotype | Estrus phase |
|---|---|---|---|
| 4362 | F | WT | estrus |
| 4341 | F | WT | estrus |
| 4325 | F | WT | metestrus |
| 4611 | F | WT | metestrus |
| 4353 | F | WT | proestrus |
| 4384 | F | WT | proestrus |
| 4381 | F | PT | estrus |
| 4373 | F | PT | estrus |
| 4380 | F | PT | metestrus |
| 4354 | F | PT | metestrus |
| 4356 | F | PT | proestrus |
| 4365 | F | PT | proestrus |

**Table S3A. Female mice from cohort generated for spine density analysis using Golgi impregnation.** A cohort comprising 6 animals per sex per genotype was generated and vaginal cytology was collected on all female animals at time of sacrifice. Groups were balanced across genotypes to include 2 animals per estrus phase from 3 phases.

| Effect | Phase | Estimate | Standard Error | DF | t Value | Pr > \|t\| |
|---|---|---|---|---|---|---|
| Cytology | estrus | 0.08222 | 0.07675 | 143 | 1.07 | 0.2859 |
| Cytology | metestrus | 0.1018 | 0.0789 | 143 | 1.29 | 0.1992 |
| Cytology | proestrus | 0.00755 | 0.07615 | 143 | 0.1 | 0.9212 |

**Table S3B. Effect of estrus phase on spine density.** A mixed model was fitted for spine density as the outcome measure with genotype, cytology, and cytology*genotype as fixed effects covariates, and animal and neuron (nested within animal) as random effects. Column G indicates the p-values for cytology in this model.

**Table S4.** Primer sequences used for qPCR.

| Target | Primers |
|---|---|
| KAL7 | F: GTTCGCAATGCACAGTGTCA<br><br>R: GCCCGTGTTAAGGTTCTCCA |
| KAL9 | F: GCCCCTCGCCAAAGCCACAGC<br><br>R: CCAGTGAGTCCCGTGGTGGGC |
| KAL9 splice | F: CCACCCAGGATGAGATGACT<br><br>R: GGTTTCTAGGAGGTGTGGGA |
| KAL12 | F: AGAACAGTGGCAATGGAGGG<br><br>R: ACCTGGGCTGCCTGAAAAAT |
| GAPDH | F: AACTTTGGCATTGTGGAAGG<br><br>R: ACACATTGGGGGTAGGAACA |

### Supplemental Methods

#### *Imaging*

Fluorophores comprising the RhoA2G sensor (mTFP and vGFP) were spectrally characterized in isolation (Nikon Inst.) and spectral unmixing (NIS Elements) was subsequently used to isolate RhoA FRET emissions (ratio of vGFP/mTFP). Neurons were usually imaged at 2Hz using a 60x water-dipping objective (NA 1.0) focused on proximal apical dendritic regions at zoom 2 (0.22 µm/pixel). mTFP was excited with 457 nm light (2%–10% laser power from a 40mW argon gas laser (note only 16mw at 457nm). The pinhole was opened slightly (1.6 airy units) to capture a broader signal and reduce sensor alterations resulting from focal changes(85).

The position of CA1 hippocampal neurons was carefully detailed at low power for subsequent imaging at higher magnification. CA1 pyramidal neurons from hippocampal slices were imaged using a 20x oil-immersion objective (NA 0.75, 0.62µm/pixel). z-stacks were collected with a 2µm step size and a maximum z-projection image was created for dendritic arbor reconstruction using NeuroLucida software (MBF Inc).

<u>Golgi</u>

For the analysis of dendritic structure *in vivo*, image acquisition was performed on a custom-built, wide-field epifluorescence Olympus IX73 microscope equipped with XYZ-encoded prior stage, real-time autofocus, and a Hamamatsu ORCA-Flash4.0 sCMOS camera controlled with SlideBook 6.0 using an Olympus PlanAPOS 40x 0.90 NA air objective. Brightfield exposure time was optimized for each image. Image stacks, 1 µM step-size, through the entire tissue depth were captured of pyramidal shaped cells within the region of interest. Pyramidal cells were included if they did not have overlapping soma from neighboring cells and their soma was located roughly within 40-60% of the z-depth. Collected image planes were 2048x2048 pixels with a resolution of 0.161µm/pixel. Fig S2A shows a representative image of a PC acquired at 40x magnification for subsequent reconstruction and analysis, and Fig S2B shows the reconstruction generated from the image. A total of 6 neurons per animal distributed across 2-3 sections were collected from the right hemisphere. For dendritic reconstruction, image stacks were manually traced offline using NeuroLucida software (Microbrightfield, Inc) and total dendritic content was analyzed via Sholl analysis using NeuroExplorer (Microbrightfield, Inc). Five animals per sex per genotype were included, for a total of 60 *Kalrn*-PT and 60 *Kalrn*-WT neurons imaged and reconstructed. Experimenter was blinded to the genotype throughout image acquisition and processing.

For the analysis of dendritic spine density, the same imaging system as described above was employed. PCs meeting the above criteria were identified at 20x magnification, and image stacks were subsequently collected using an Olympus PlanAPOS 60x 1.42 NA oil objective at a step size of 0.7 µm. For spine counting, image stacks were manually evaluated offline using StereoInvestigator software (MBF Bioscience). A portion of the apical dendrite spanning 20-150 µM from the soma was demarcated (Fig 5A), and any secondary apical dendrite emanating from that region was including for spine counting if it had at least 10µm of length before branching or terminating. A neuron needed to contain at least 100µm of secondary apical length for spine counting to be included in the final analysis. Three to five neurons per animal were included in the final analyses.

<u>Nissl</u>

For cortical volume estimates, imaging was performed on an Olympus BX51 WI upright microscope (Olympus). Images were captured at 1.25x.

<u>IHC</u>

Imaging was performed using a 60X oil supercorrected objective (NA 1.42) mounted on an Olympus BX51WI upright microscope equipped with an Olympus DSU spinning disk, Hamamatsu Orca R2 camera, MBF CX9000 front mounted digital camera (MicroBrightField Inc.), BioPrecision2 XYZ motorized stage with linear XYZ encoders (Ludl Electronic Products Ltd.), excitation and emission filter wheels (Ludl Electronic Products Ltd.), Sedat Quad 89000 filter set (Chroma Technology Corp.), and a Lumen 220 metal halide lamp (Prior Scientific). The process of image collection was accomplished using Slidebook software version 6.0 (Intelligent Imaging Innovations) and Stereo Investigator version 8 (MicroBrightField Inc.). Detailed description of image acquisition and analysis is provided in (9). Briefly, images were processed using SlideBook version 6.0 (Intelligent Imaging Innovations) with keystrokes automated by Automation Anywhere software (Automation Anywhere, Inc). Image stacks were deconvolved using the AutoQuant adaptive blind deconvolution algorithm (Media Cybernetics). Spinophilin-IR and phalloidin puncta with intensity measures above the Ridler Calvard defined minimum threshold derived value in SlideBook were selected, and contiguous pixels were defined as a "mask object." Identification of putative dendritic spines required colocalization of spinophilin-IR and phalloidin label, defined as phalloidin mask objects that overlapped (≥1 voxel) with a spinophilin-IR mask object. Spine density ($N_v$) and number ($N$) in cohort 1 were calculated as previously described (9).

#### ***CRISPR/Cas9***

<u>Mouse Production</u>

We chose a single male founder that was homozygous for the knockin mutation to establish the line. He was mated with a wildtype C57BL/6J female purchased from Jackson Laboratories (Bar Harbor, ME) and all F1 mice were genotyped by both PCR and restriction digest as well as confirmatory Sanger sequencing to establish the *Kalrn-PT* line. Heterozygotes were backcrossed into purchased wildtype C57BL/6J strain to produce heterozygous breeders at each generation to produce animals for all experimental studies mitigating against genetic drift of the line.

Animals were identified with metal ear tags and tail snip DNA samples were obtained for genotyping by PCR prior to weaning at approximately postnatal day 21. Genotypes of all mice included in studies were confirmed following sacrifice. Both male and female animals were included in all studies. Vaginal cytology was obtained on all adult female mice on the day of sacrifice to determine phase of the estrus cycle. Animals were under specific pathogen free conditions and housed in standard microisolator caging (Allentown Caging Equipment, Allentown, NJ, USA) in groups of up to 4 males or 5 females, maintained on a 12-h light/dark cycle (lights on at 7 AM), and were provided with food and water ad libitum. All experimental procedures were approved by the Institutional Animal Care and Use Committee at the University of Pittsburgh.

Studies were performed on wildtype and homozygous knockin mice, which for simplicity are referred to throughout the text as *Kalrn-WT* and *Kalrn-PT*.

#### ***Golgi***

In brief, the extracted brain was quickly rinsed with distilled and deionized water and placed into a Golgi-Cox solution composed of potassium dichromate, mercuric chloride, and potassium chromate. The Golgi-Cox solution was changed the following day to fresh solution, and the brains were incubated in the dark at room temperature with gentle agitation for 2 weeks. The brains were then transferred to a cryoprotectant solution and incubated in the dark for 3-6 days. Brains were sectioned (150 µm) on a cryostat. Tissue sections were placed onto gelatin-coated slides and allowed to dry overnight at room temperature. The following day sections were then rehydrated with distilled and deionized water, reacted in a developing solution (FD Rapid GolgiStain Kit, FD Neurotechnologies), and dehydrated through graded ethanols. Finally, sections were cleared in xylene and coverslipped using Permount (Fisher Scientific). Age-matched cohorts that included males and females of both genotypes were processed together throughout.

***Immunohistochemistry (IHC)***

IHC was carried out as previously described (9). In brief, free-floating sections were pre-treated with 1% NaBH4 to reduce auto-fluorescence and 0.3% Triton X in order to permeabilize the sections before being incubated for two hours at room temperature in blocking buffer (20% normal goat serum, 1% bovine serum albumin, 0.1% lysine, 0.1% glycine, and phosphate buffered saline (PBS)). Following blocking, tissue was immediately incubated with primary antibody overnight at 4˚C. Sections were rinsed in PBS then incubated with Alexa Fluor® 488 goat anti-rabbit (1:500; Invitrogen A11034) and Phalloidin- Alexa Fluor® 568 (3:200) at 4˚C for 24 hours. The tissue sections were subsequently mounted on gel coated slides, rehydrated to ameliorate the effects of z-axis tissue shrinkage, and mounted onto gel coated coverslips using Vectashield hard-set H-1400 mounting medium (Vector Laboratories).

***Determination of cortical volume***

A subset of an additional cohort of 12 week ± 3 days mice (see IHC previously, 8F and 12M, 10 animals per genotype) was used. Upon sectioning, every 8$^{th}$ section was pulled for Nissl staining, providing a fixed interval of 320µM between Nissl-stained sections for each animal. Using a random start of either section 8 or 16, every other Nissl section (providing a fixed interval of 640µM between sections) was included for a given animal for the full rostral-caudal extent. Using the mouse atlas as reference, contours were drawn outlining the neocortex (see Fig S4). Approximately 8-12 sections per animal were contoured and imaged.

Estimates of cortical volume were determined using the Cavalieri method (92). A square point grid with an *a/p* of 1.6mm$^2$ was superimposed at a randomized angle on each section, and the volume of the neocortex ($V_{cortex}$) was calculated for each animal as: $V_{coretx} = T \times (a/p) \times \sum P$ where ∑P is the number of points counted per region and T is the intersection distance (0.64mm, as every other Nissl section was counted). Mean (SD) points counted per animal was 410.9 (50.8).
